# Flower color locus resists introgression due to correlational selection with other floral traits in *Ipomoea cordatotriloba*

**DOI:** 10.64898/2026.04.29.721739

**Authors:** Jonathan Z Colen, Mark D Rausher

## Abstract

- When species hybridize, resistance to introgression is presumably due to selection against hybridizing alleles. While many studies have characterized direct selection at these sites, alleles may resist introgression through correlational selection. Here we investigate the role of direct and correlational selection in reducing introgression at the color locus in *Ipomoea cordatotriloba*.
- We used recombinant inbred lines that varied in limb color, flower size and sugar concentration to estimate the fitness advantage of the flower color allele via direct and correlational selection. To assess the effect of correlational selection on fitness, we ask if floral size or nectar sugar concentration is correlated with fecundity in pink- but not white-limbed lines.
- We find no evidence for direct selection on flower color across four fitness components – germination, survival, fecundity, and siring success. Instead, both flower size and sugar concentration significantly correlate with fecundity in pink, but not white limbed lines. As a result, correlational selection on the color allele opposes introgression when recurrent migration is low (<3%).
- These results demonstrate that correlational, rather than direct, selection is sufficient to resist introgression via hybridization and suggest that correlational selection is an underexplored mechanism to generate resistance to introgression across multiple loci.

## Introduction

Since Darwin (1859), evolutionary biologists have puzzled over a simple question: how do species remain distinct when they readily hybridize? Interspecific mating is quite common in the natural world – over a quarter of all plant species readily hybridize (Mallet 2005). One potential consequence of hybridization is introgression, defined by Rieseberg and Wendell (1993) as "the permanent incorporation of genes from one set of differentiated populations into another". The extent of introgression is determined by two factors: the strength and pattern of selection on introgressing alleles, and the rate of gene flow between populations. Depending on these factors, several different outcomes are possible. First, if introgressed variants are advantageous across populations, then they may be universally fixed (Suarez-Gonzalez et al. 2018, Taylor and Larson 2019). Most alleles that introgress, however, are presumably neutral. In this case, gene flow from one population into the other provides a continual input of the introgressing allele. If this gene flow is recurrent and asymmetric (i.e. more frequently from population A to population B than the reverse), neutral alleles from the donor population (A) will eventually be driven to fixation in the receiving population (B).

A final possible outcome is resistance to introgression despite recurrent gene flow. If there is purifying selection against the introgressing allele, then that allele will increase only to low or intermediate frequency, with the allele’s final frequency reflecting the balance between the strength of purifying selection and the magnitude of gene flow. When purifying selection is weaker than the strength of gene flow, introgressing alleles are effectively neutral and will become fixed due to recurrent gene flow. By contrast, if purifying selection is strong and the magnitude of gene flow is low, selection can effectively prevent any introgression (for further elaboration, see model results below).

Recent advances in genomic sequencing (see Seehausen et al. 2014) have shown that genomes appear as “mosaics” (Payseur 2010), with some regions that readily introgress and others that show little to no introgression (Wu 2001, Payseur and Rieseberg 2016). Loci with high admixture presumably reflect either adaptive introgression or sufficient gene flow to drive neutral alleles to fixation. By contrast, loci with little to no sign of introgression (i.e., low admixture) presumably experience purifying selection opposing gene flow. To substantiate this inference, researchers often sequence hybridizing populations, identify candidate loci that resist introgression, and posit that this lack of inrogression is caused by purifying selection (e.g. Payseur 2010). However, these genomic analyses are not sufficient for inferring definitively whether purifying selection has operated, as there is no direct test of selection. Moreover, these analyses cannot elucidate causal mechanisms responsible for selection (Suarez-Gonzalez et al. 2018, Taylor and Larson 2019). In particular, when multiple traits resist introgression, selection may either act independently across all traits or jointly in the form of correlated selection. Despite extensive evidence that traits jointly resist introgression (Rieseberg et al. 1999, Fishman et al. 2014, Schumer et al. 2014, Rifkin et al. 2021, Hankin 2024), there are only two studies that explicitly test whether these traits are jointly or independently selected (Meléndez-Ackerman and Campbell 1998, Campbell 2009).

Three mechanisms could generate resistance to introgression. First, the introgressing variant could be directly selected against, either because the phenotype it produces is maladaptive, or because the variant has deleterious pleiotropic effects on another trait. Second, it could be selected against because the focal trait is genetically correlated with one or more other traits that experience purifying selections (Lande and Arnold 1983, Hawthorne and Via 2001, WHEAT et al. 2011, Comeault et al. 2015, Friedman et al. 2015); theory suggests that even moderate genetic correlations between traits can cause resistance to introgression (Lindtke and Buerkle 2015). Third, resistance to introgression could be caused by correlational selection involving the trait of interest and other traits. In this case, selection on the focal trait is conditional on the phenotype of other traits even when genetic correlations between these traits are low (see below for further explanation). This process differs from the second mechanism in that correlational selection is a pattern of selection, while genetic correlations cause an indirect response to selection that does not require correlational selection.

To our knowledge, the possibility that correlational selection can cause resistance to introgression has not been previously considered. Experimental studies that quantify selection against introgression are uncommon, and nearly all such studies attempt to detect only direct selection on the focal trait (e.g. Bradshaw et al. 1998, Fulton and Hodges 1999, Hopkins and Rausher 2012, Kenney and Sweigart 2016, Farnitano et al. 2025). Because many loci typically resist introgression when species hybridize (Rieseberg et al. 1999, Fishman et al. 2014, Schumer et al. 2014, Rifkin et al. 2021, Hankin 2024), it is surprising that the contribution of trait correlations have seldom been investigated as possible mechanisms underlying resistance to introgression. We are aware of only one such system, which involves introgression between *Ipomopsis aggregata* and *I. tenuituba* (Meléndez-Ackerman and Campbell 1998, Campbell 2009). Consequently, we have yet to understand the general importance of correlational selection at loci resisting introgression.

Correlational selection is a form of fitness epistasis: the value of one trait (focal trait) affects the magnitude or direction of selection a second trait. In this study, we consider the case where the focal trait is controlled by a single locus, which modifies the pattern of selection on a second, quantitative trait. As we show below, if this modification changes the direction of selection on the quantitative trait, the resulting feedback can generate resistance to introgression at the focal trait locus. Notably, this mechanism is distinct from epistatic mechanisms contributing to speciation, e.g. epistatic selection on hybrid incompatibilities (Gavrilets and Hastings 1996, Gavrilets 1997), and in generating assortative mating (Felsenstein 1981, Barton and De Cara 2009), since these processes involve two Mendelian traits rather than a quantitative trait.

We test the extent to which correlational and direct selection contribute to resistance to introgression in a system with recurrent, asymmetric gene. *Ipomoea lacunosa* and *I. cordatotriloba* are two sister species that separated about 1 million years ago (Carruthers et al. 2020), and have subsequently come into secondary contact across the coastal plains of the southeastern United States. *I. cordatotriloba* has a mixed mating system (Mc Donald et al. 2011), and where both species co-occur, it can be pollinated by *I. lacunosa* and form fertile hybrids, despite strong incompatibility barriers (Ostevik et al. 2021, Rifkin et al. 2023). In contrast, *I. lacunosa* is highly selfing (selfing rate >95%; Duncan and Rausher 2013), and there is little opportunity to produce hybrid seeds. Because of this asymmetry, hybridization unidirectionally introduces *I. lacunosa* alleles into sympatric *I. cordatotriloba* populations (which we define as “introgression”). These processes result in two sympatric populations that are largely, though not completely, reproductively isolated (Ostevik et al. 2021): “sympatric *I. cordatotriloba*” that is phenotypically similar to *I. cordatotriloba* with high levels of introgression from *I. lacunosa* (average admixture = 62%), and *“*sympatri*c I. lacunosa*” that is phenotypically indistinguishable from *I. lacunosa* with virtually no introgression from *I. cordatotriloba* (average admixture < 2%) (Rifkin et al. 2019a).

The extent of introgression into *I. cordatotriloba* varies across the genome. At many loci, introgression has led to fixation or near fixation of *I. lacunosa* alleles. By contrast, other loci effectively resist introgression and exhibit little admixture (Rifkin et al. 2019b), presumably because purifying selection acts against the *I. lacunosa* allele at those loci. One such locus is a flower color locus. In allopatry, nearly all *I. lacunosa* flowers have white limbs (Stephenson IV et al. 2006) and *I. cordatotriloba* flowers have pink limbs (Duncan 2013), with flower color difference controlled by a single Mendelian locus (Duncan and Rausher 2020; see Materials and Methods). In sympatry, almost all *I. cordatotriloba* flowers also have pink limbs, with white-limb color occurring only at low frequency (mean p = 0.078) (Duncan 2013). *I. cordatotriloba* thus appears to be resistant to introgression at this locus. We explicitly tested the hypothesis that selection generates this resistance by quantifying direct and correlational selection on the flower color locus in *I. cordatotriloba*.

To quantify direct selection on flower, we estimated four components of selection at the flower color locus in experimental populations: germination proportion, survival to flowering, seed production, and paternal pollination (siring) success. To specifically determine whether correlational selection contributes to resistance to introgression at the color locus, we focus on two important floral traits - flower size and nectary sugar concentration - that, along with flower color, commonly affect pollinator visitation in plants (Conner and Rush 1996, Fowler et al. 2016, Fornoff et al. 2017, Mallinger and Prasifka 2017, Nottebrock et al. 2017, Delgado et al. 2023). In accordance with the model of correlational selection presented below, we determined whether the pattern of selection acting on these two traits depends on flower color and generates purifying selection against the allele for white limb color.

## Materials and Methods

### RIL creation and selection

We used recombinant inbred lines (RILs) because they allowed estimation of fitness of the two homozygote genotypes at the flower-color locus while randomizing much of the genetic background. RILs were previously generated from a cross between a pink-limbed sympatric *I. cordatotriloba* (CAA) and a white-limbed *I. lacunosa* (LPR; see Rifkin et al. 2019b for cross details). Hybrid offspring were selfed for five generations to produce approximately 300 fifth (F5) and sixth (F6) generation RILs. For these lines, floral size, color, and nectar traits were quantified (see Liao et al. 2022). Because genetic correlations that originally existed in sympatric *I. cordatotriloba* were disrupted via recombination during the generation of the RILs, the independent effects of color, flower size, and sugar concentration can be individually quantified. 118 lines only had pink or white limbed color across generations and were considered to be homozygous for the respective color allele – 59 lines for each limb color. Because flower size traits are highly correlated with each other, as are nectar traits (see Liao et al. 2022), we chose one representative trait – corolla length and nectar sugar concentration – from each category for analysis. These traits were quantified for the 6^th^ generation inbred lines in a previous experiment (Liao et al. 2022). Separately, we confirmed that limb color segregates in Mendelian fashion. Allopatric *I. cordatotriloba* was crossed with *I. cordatotriloba* homozygous for the white allele, found at low rates in sympatric populations (see Duncan 2013). Among F2 offspring, 166 were pink-limbed and 56 white-limbed, producing a ratio of 2.96:1, which is consistent with a 3:1 ratio (p-value=0.938, G-test=0.00594).

### Field experiment design

In June 2021, 25 *I. lacunosa*, 25-white-limbed RILs, and 25-pink-limbed RILs were planted in 16 replicate plots (4×4 grid) at a common garden at the Border Belt Tobacco Research Station (BBTRS) in Whitesville, NC, 11.7 kilometers from the closest known sympatric population in Chadbourn, NC (34.31021N, −78.82640W; CHAD in Rifkin et al 2019b) and 63.7 kilometers from the closest known allopatric population in Supply, NC (34.01894N, −78.30006W; Site 3 in Rifkin et al 2019b). Plots were arranged in a randomized block design, with each plot containing one plant per RIL (50 RIL plants) and 25 *I. lacunosa* randomized throughout the plot. Each replicate plot used the same 25 RILs for each color type, except for two white-limbed RILs that did not have enough seeds to plot sixteen replicates; for these two lines, a separate white-limbed RIL was used for half of the replicate plots. *I. lacunosa* was planted both to assess micro-environmental quality (as “phytometers”; see below) and to quantify any fitness cost of pollen deposition onto *I. cordatotriloba* stigmas. Plants were spaced at 1-meter intervals in a square grid, with no plant directly bordering another plant of the same type (Figure S1).

Seeds were scarified, planted in trays, and, upon producing their first true leaf, were transplanted into slightly raised ground between furrows. Plants were initially irrigated to prevent transplant shock; five plants died within the first two weeks and were replaced. Survival was then scored at least once a week until a plant produced its first flower. Three weeks post-transplant (July 2), two metrics of early growth were recorded: (1) the total number of shoots, and (2) the length of the tallest shoot, measured from the base of the plant to the tip for the longest living vine. Because the two traits are correlated (p<2.2*10^-16^, R^2^=0.4134), only the results for shoot length are discussed.

### Germination Assay

To assess whether white- and pink-limbed RILs differed in germination probability, 8000 full-sib seeds were planted at the BBTRS in June 2022. Soil was sifted into 1” PVC pipes sunk into the ground to prevent contamination from the local seed bank. For 10 pink-and 10 white-limbed RILs, 400 seeds were planted per RIL. Ten seeds from each RIL were planted in each of 40 replicate pipe containers. Containers with pink-limbed seeds never directly bordered containers with white-limbed seeds, but RIL ID was randomized within color type. Every 4-7 days, we counted the cumulative number of new germinants since the last observation period until the final census on October 20^th^. Once cotyledons were unfurled, a plant was removed to prevent shading of nearby seeds. We calculated the germination proportion per line by dividing the number of germinants by 400, the number of seeds planted per line.

### Assessing Female Fitness

To estimate whether pink- and white-limbed lines differ in seed production, seeds were collected from each plant by collecting senesced plants and storing them in paper bags. Because plants senesce shortly after peak flower production and seeds remain in their capsules for long periods after maturation, seeds could be accurately collected and counted. Subsequently, seeds were manually separated from dead plant tissue and collected in coin envelopes for each plant. Because of the time-intensive nature of seed separation, seeds were extracted for only four adjacent plots of the sixteen original plots.

All seeds from each plant were weighed in bulk. To convert seed mass to seed number, fifty haphazardly chosen seeds were weighed to determine the average mass of a seed per sample, and the total mass was then multiplied by the estimated number of seeds per gram. To validate the accuracy of these estimates, 24 randomly sampled individuals across four plots were hand-counted and compared to estimated seed numbers; estimated seed number was highly correlated with counted seed number (Fig S2, p<2*10^-16^, R^2^=0.9883). For nearly all samples (22 of 24 samples), actual seed number was within 10.9% of estimates (Figure S2), with the two outlier plants having substantial seed predation post-harvest from burrowed stinkbug instars not previously detected when seeds were sorted and weighed. Because both viable and inviable seeds were collected together, one plot (Plot A) was selected to test whether the proportion of inviable seeds differed among flower color treatments. Viable and inviable seeds were manually separated and weighed to estimate the proportion of seed mass which was inviable. No significant differences in the fraction of inviable seeds were detected across color types (p= 0.9582; Table S1). Because seed number and seed mass are highly correlated (p<2*10^-16^, R^2^=0.9455), analyses using line estimates of seed number produce qualitatively similar results compared to those using seed mass.

### Using I. lacunosa as Phytometers

Typically, plants growing in good micro-environmental conditions will be larger and set more seeds than plants growing in poorer conditions. To remove the effect of local environment on plant size and on seed set, we use the early plant size of *I. lacunosa* as a phytometer - a proxy for assessing local environmental quality (Clements and Goldsmith 1924). With this proxy, we corrected seed set estimates of RIL plants using two different models – a polynomial model (Model 1) and a spline-based model (Model 2; see supplementary methods for more information).

### Assessing Siring Success via Sanger Sequencing

To assess the relative siring success of pink- and white-limbed RILs, we used a marker-based paternity analysis similar to those we have used previously in other studies (Rausher et al. 1993, Rausher and Fry 1993, Fehr and Rausher 2004). Because F5 RILs were previously RAD sequenced (Liao et al. 2022), we conducted a genome-wide scan of all sites to identify sites where one marker allele was above 90% frequency in all pink-limbed lines, and an alternative allele was above 90% frequency in all white-limbed lines. There was only one sequenced site that met this criterion (15210053 on Chromosome 6 (Table S2)). At this site, most pink-limbed RILs were homozygous for nucleotide A, while most white-limbed RILs were homozygous for G. We confirmed this pattern using PCR of each RIL. We extracted leaf tissue using a slightly modified version of the protocol of Murray and Thompson (1980), in which ethanol washes replace salts and other forms of alcohols to precipitate DNA. For PCR, we amplified our target sequence by combining genomic DNA with one primer ∼170 bp downstream of the target site (sequence: TGTAGGGTTGAGGAGAGAGGT) and the other ∼50 bp upstream of the target site (sequence: AGAAGGAGATGGGTGAAGGG) using an Biometra TOne Thermal Cycler (Analytik Jena). The PCR ran for 33 cycles at 95C/52C/72C.

After verifying that this marker allele is strongly associated with color for the 53 RILs used for estimating siring success, we randomly sampled 200 offspring seeds from each of three plots, weighting the probability of sampling a given plant by its seed production. PCR products from each offspring were Sanger sequenced by Eurofins Genomics. Because of germination failure, unsuccessful DNA extraction, or poor sequence quality, genotype could be determined for only 393 samples. To ensure that these samples (194 from pink-limbed and 178 from white-limbed mothers) were reflective of the original 600 sampled seeds, we asked if a) the proportion of seeds from pink- vs white-limbed lines, and b) proportional contributions from each line, were statistically indistinguishable using a Chi-square test. In all cases, we found no statistical differences (Table S3), suggesting loss of samples did not create bias. Samples were scored as homozygous or heterozygous by analyzing chromatographs in Snap Gene Viewer (see Figure S3).

### Likelihood Model of Siring Success

To estimate the proportions of pink-limbed plants sired by either white-limbed or pink-limbed fathers, and the proportions of white-limbed plants sired by either pink-limbed or white-limbed fathers, we employed a likelihood analysis. In the Supplementary Material we show that the per-capita rates of siring success for pink-limbed and white-limbed parents, 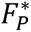 and 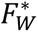, respectively, are given by dividing Eqs. (1a) and (1b) by the respective number of potential sires:

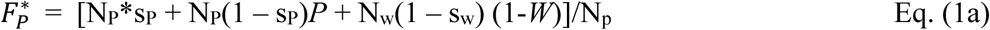

and

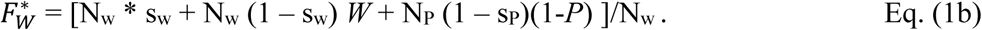

where N_P_ and N_W_ are the numbers of pink- and white-limbed plants, sp and sw are the corresponding selfing rates, *P* is the probability that a pink-limbed plant is pollinated by pink pollen, and *W* is the probability that a white-limbed plant is pollinated by white pollen. We also show that, when N_p_ = N_w_, as in our experiment, and s_p_ = s_w_, as we show in the Results section, a test of *P* = *W* is equivalent to a test of whether 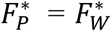, i.e. whether pink- and white-limbed plants have equal *per-capita* siring success.

### Statistical Analysis for Relationships among Traits

Across three of the four measured fitness components (germination rate, survival, seed set), differences between pink- and white-limbed lines were calculated using type II ANOVA with color as the independent categorical variable, each metric of fitness as the dependent variable, and an aspect of location of each seedling or plant as random effect covariates. Because phytometer values did not effectively control for local environmental variation (see Results), location of plants is considered as a random effect. In the analysis of germination, this location is the row and column coordinates of the different containers that seeds were planted in, and for survival, early growth, and seed set, this random effect is block. When line averages are used, the average row and column coordinates are used as random effects. For any analysis not conducted on line means, the effect of RIL was also treated as a random effect. To quantify relationships between color, flower size, and sugar concentration, we employed generalized linear mixed models with plot and line as random effects using the lme4 (v1.1.36, Bates et al. 2015) and car (v. 3.1.3, Fox and Weisberg 2018) packages in R. Generalized linear mixed models were of the binomial family for models with survival as the outcome variable and of the Gaussian family when germination, early growth and seed set were the outcome variables.

Models included flower size, sugar concentration, and color as fixed effects, and, when relevant, included position in plot as a random effect. Because Type II and Type III ANOVAs test the same parameters when there are no interacting effects (Langsrud 2003), significance was calculated using type II ANOVA for direct effects of color, sugar concentration, and flower size, and type III ANOVA for interactive effects via the car package. All analyses were conducted using version 4.4.0 in R (R Core Team 2024).

### Modelling Maintenance of Flower Color

To examine the hypothesis that selection against the white allele was generated by correlational selection, we constructed a mathematical model, guided by results from the field experiment. We assumed that the fitness of a pink plant is *W_p_* = α + *β_P_*\*z, where z is the value of the quantitative trait for that plant. By contrast, we assumed that *W_w_*, the fitness of a white plant, is

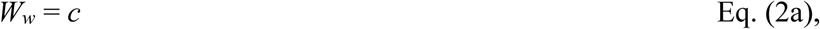

i.e. constant and not affected by z. The average fitness of a pink plant can be re-written as

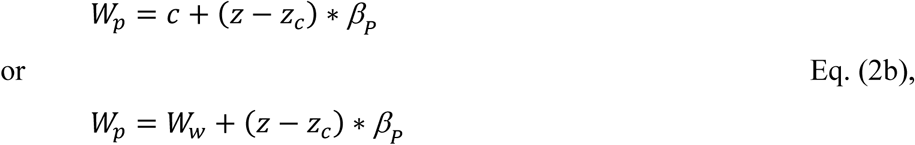

where *z_c_* is the trait value at which *W*_*p*_ = *W*_*w*_.

We used Equations 2a,b to construct an explicitly genetic model in which color phenotype determined by genotype at a single “color” locus, and a value of the quantitative trait determined additively by the effects of two alleles at each of k loci. We assume no linkage among any of the loci. One allele, the “+” allele, increases the trait by a fixed increment, δz, while the “-” allele decreases the trait by the same fixed increment (Figure 1). We implemented two different versions of the model to control for programming error: one constructed in Mathematica and one constructed in R. Both provide the same qualitative output (see Supplementary Methods). Each modeling implementation is comprised of 5 components. First, we initialized a population to be fixed for both the pink color allele and for the “+” allele at *k* quantitative trait loci. Second, we implement immigration at rate *m* of gametes that carry the white color allele and “−” quantitative trait alleles. Third, we calculate the resulting genotype frequencies, assuming random mating between gametes. Fourth, we impose selection of genotypes, where the fitness of each genotype is calculated based on Eqs. 2a,b. Lastly, for each genotype, we generate all possible gametes with frequency (½)^j^, where j is the number of heterozygous loci. These gametes contribution to the next generation’s gamete pool is proportional with genotype frequencies. Details are presented in the Supplementary Methods.

**Figure 1.**
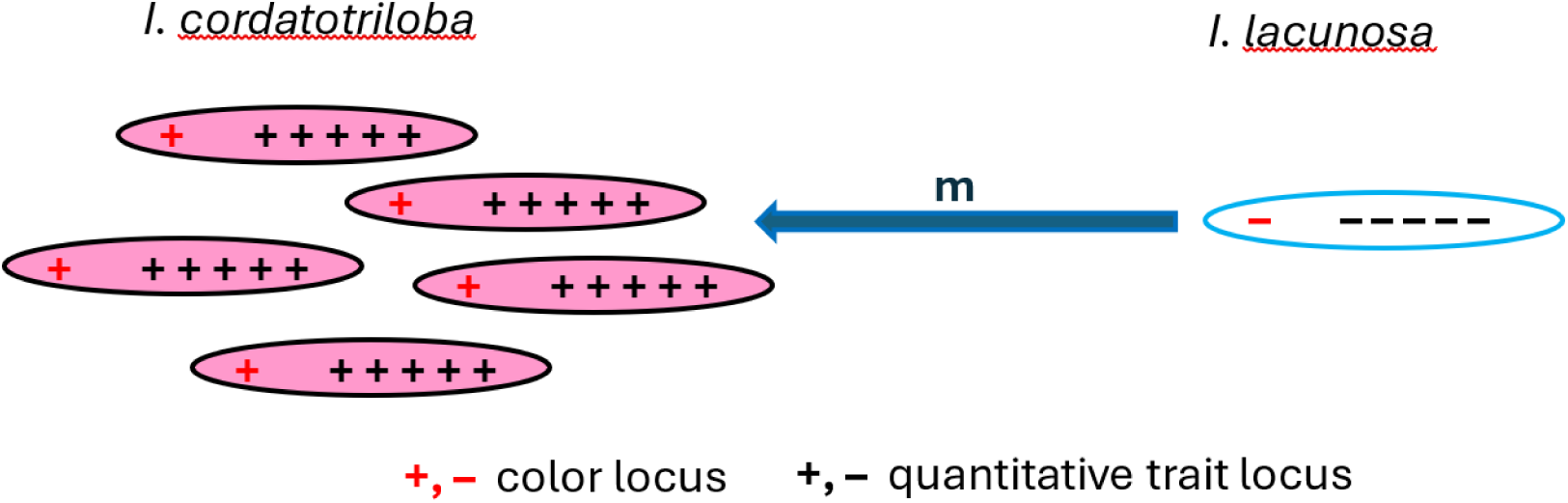
Migration of *I. lacunosa* haplotype into sympatric *I. cordatotriloba*. Populations start at 100% frequency of pink allele (red “+”) and alleles responsible for an increased quantitative trait (black “+”) alleles. Each generation, migration adds the white allele (red “-”) and alleles for a decreased quantitative trait (black “-”) at rate *m*. This process occurs recursively for 500 generations after selection and gamete formation.

Parameters for the model were estimated from the experimental data. For *β_P_* we used the slope of the regression of seed mass on flower size, 1.878. For *z_c_* we used the flower size at which finesses of pink and white-limbed lines are equivalent, 26.3. For *c* we used the mean fitness of white-limbed plants, 20.4. Finally, for *z* we assumed that the extreme values of flower size (21.562mm and 31.038mm) represented individuals with either 10 “+” or 10 “−” alleles. The incremental change in *z* due to adding 1 “+” allele was then assumed to be used 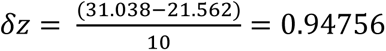; therefore *z* = 21.562 + *n δz*, where *n* is the number of “+” alleles an individual carries.

### Linkage Disequilibrium Analysis

Lastly, we used our model to evaluate the role that correlated selection has in increasing linkage disequilibrium (LD) between the color locus and quantitative trait loci. LD was measured as the difference between haplotype frequencies and their expected frequencies given the product of each gamete frequency (Slatkin 2008). To assess the proportion of additive genetic variance explained by LD, we calculated the correlation coefficient (r_G_) between the color locus and phenotype, such that a value of r_G_ of 1 would indicate that quantitative traits co-associate with the color locus and are in complete LD, and a value of 0 would indicate no associations between color loci and quantitative trait loci. To quantify the proportion of allele frequency change due to correlational selection as compared to direct selection on traits, we first calculated the allele frequency changes due to selection as done above. We then constructed a model that artificially removes linkage disequilibrium by setting initial gamete frequencies at their expected values from each allele frequency. Differences across these models reflect the change in allele frequency caused by LD. For both models, we arbitrarily chose a value of *m* (0.04) for which the allele would remain at migration-selection balance and evaluated the change in frequency of the pink allele due to selection.

## Results

### No Direct Selection on Flower Color

Lifetime fitness in an annual plant can be partitioned into several multiplicative components:

W_lifetime_ = Prob[germination] * Prob[survival to flowering] * ½ [*N_seeds_* + *N_pollen_*], where *N_seeds_* is the number of seeds produced and *N_pollen_* is the number of pollen grains produced that sire ovules. Here we examine these fitness components and ask whether they differ between pink- and white-limbed plants.

#### Germination

Germination was uniformly low across all lines, with an overall average germination rate <0.11 (Figure S4). Color has no detectable effect on germination rate (binomial generalized linear model (***χ***^**2**^= 0.1699; p=0.6802, Figure **2a**). Because germination time may potentially affect fitness, we also asked whether pink- and white-limbed lines differed in mean germination time. While pink-limbed lines germinate earlier, and this difference is significant (***χ***^**2**^=4.6408, p=0.03122), the time difference is less than 4.5 days (Figure S5), suggesting that this difference in germination time does not substantially impact fitness.

**Figure 2.**
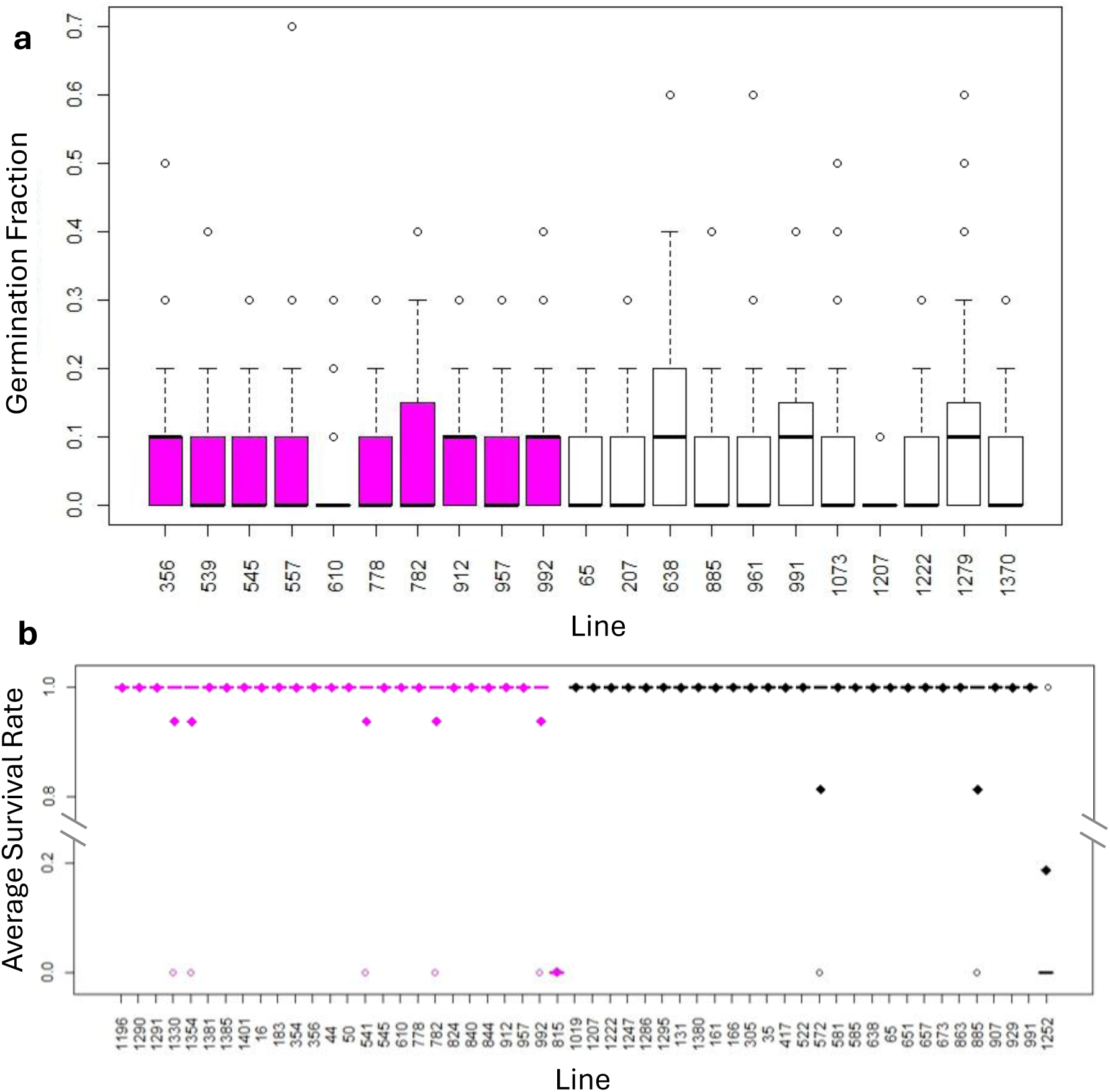
No significant differences in germination rate nor survival between pink- (pink bars) and white- (white bars) limbed lines. **a)** Germination proportion for pink- (shaded pink) and white-limbed (shaded white) lines. Black lines represent averages for each color, whiskers represent 1.5 times the interquartile range (IQR), boxes the middle 50% of data range. Circles represent data points outside of this whisker range (1.5 *IQR). **b)** Survival to first flower for pink-(pink-colored) and white-limbed (black colored) lines. Diamonds represent line averages, dashes represent whether the median plant per line lived or died, and open circles represent outlier plants that survived or died. No lines had an average survival rate between 0.2 and 0.8.

#### Survival to flowering and plant size

In contrast to germination, survival to first flower was uniformly high across all RILs and for control *I. lacunosa*, with survival rates >94.5% across all categories (Figure **2b**, Table S4). To test for differences in survival between pink- and white-limbed lines, we performed a binomial generalized linear mixed model analysis, with color as the fixed dependent variable with random effects of RIL line and block (see Methods). Differences in survival were highly line specific; two lines (one pink-limbed and one white-limbed) had a survival rate below 20%, seven had a survival rate between 80% and 94%, and the other 45 lines had a 100% survival rate. We find no significant difference in survival to first flower between pink- and white-limbed lines (survival rate for pink- and white-limbed lines is 0.9495 and 0.9575, respectively; SE=0.0383, 0.0300, respectively; ***χ***^**2**^=0.0232, p=0.8788 via ANOVA; Figure **2b**). In addition to survival, we also quantified plant growth rate (measured as plant size at two weeks). If plants are larger at flowering, and that discrepancy in size at flowering is due to the rate of early growth, then plants that grow faster early in the season may produce more flowers and thus increase subsequent fitness components (seed and pollen production). However, pink- and white-limbed plants showed no significant differences in early growth rate (size at 2 weeks) after accounting for effects of block and RIL (Figure S6, ***χ***^**2**^=0.5363, p=0.464 via ANOVA).

#### Seed Production

When uncorrected for spatial environmental variation, neither total seed mass (Figure **3a**, ***χ***^**2**^=0.9091; p=0.3404) nor estimated seed number (Figure **3b**, ***χ***^**2**^=1.3527, p=0.2448) differed significantly between pink- and white-limbed lines. However, block is a crude indicator of the local environmental variation that may affect seed production. Because early growth is a significant predictor of seed set in these plots (p=2.855*10^-5^, R^2^=0.08433; Figure S7, Table S5) and plant growth is commonly believed to be a reliable proxy for environmental quality (Dietrich et al. 2013), we used the early growth of *I. lacunosa* as a phytometer – i.e. an indicator of fine-scale variation in environmental quality – to better control for the effects of local environmental quality (see Materials and Methods).

**Figure 3.**
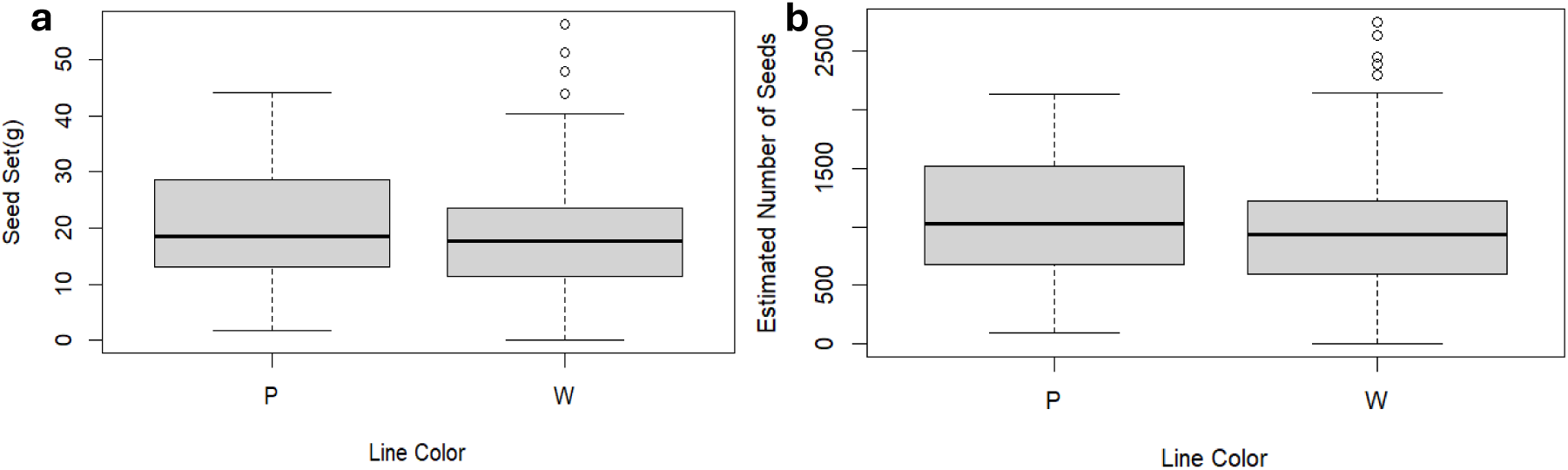
No significant differences in seed production between pink- and white-limbed lines. **a)** Mass of seed set per plant, in grams, for pink (P) and white-limbed (W) plants. **b)** Estimated number of seeds per plant for pink (P) and white-limbed (W) plants. Black lines represent averages for each color, whiskers represent 1.5 times the interquartile range (IQR), grey boxes the middle 50% of data range. Circles represent data points outside of this whisker range (1.5 *IQR).

We employed two models for phytometer effects, each of which estimates the relationship between environmental quality and position in the experimental grid. Model 1 fits environmental quality as a quartic regression surface, while Model 2 fits environmental quality as a spline estimated surface (see Materials and Methods). For both models, there is no correlation between phytometer values and seed set (Table S5, Fig S8; Model 1: p=0.2495, adj R^2^=0.004623; Model 2: p=0.8727). Moreover, for neither model did adjusted seed mass differ significantly between pink- and white-limbed lines (p=0.433 for polynomial surface; p=0.1156 for spline, Table S6). It thus appears that accounting for local micro-environmental variation does not alter our conclusion that there is no evidence that flower color affects seed production.

#### Siring Success

Approximately 80% of seeds analyzed are consistent with intra-color mating (including selfing) (Table 2). By contrast, approximately 17% of matings were between colors, with an additional approximately 13% of matings being ambiguous (Table 1). To evaluate whether these inter-color matings reflect differences in siring success, we employed a series of likelihood models with increasing constraints (Table 2) to test if pink-limbed plants have greater siring success than white-limbed plants. We first tested the null hypothesis that the proportions of alleles *A* and *G* in pollen from pink and white plants, respectively, equal their proportions among the RILs (i.e. *a =g*) by comparing Model 1 (with these proportions fixed to their actual values) with Model 0 (all parameters unconstrained). Because this constraint did not significantly decrease the log likelihood (Table 3), we fail to reject the null hypothesis and include this constraint in subsequent models.

**Table 1.** Number of offspring genotypes for given parental genotype and flower color.

| *Plant Category (Parent genotype -> offspring)* | *Number of offspring* | *Proportion* | *Mating Type* |
| --- | --- | --- | --- |
| $N_{PAA \rightarrow AA}$ | 160 | 0.413 | Within color |
| $N_{PAA \rightarrow AG}$ | 35 | 0.090 | Between colors |
| $N_{PGG \rightarrow AG}$ | 10 | 0.026 | Between colors |
| $N_{PGG \rightarrow GG}$ | 3 | 0.0078 | Within color |
| $N_{WAA \rightarrow AA}$ | 0 | 0 | Within color |
| $N_{WAA \rightarrow AG}$ | 0 | 0 | Between colors |
| $N_{WAG \rightarrow AA}$ | 5 | 0.129 | Either |
| $N_{WAG \rightarrow AG}$ | 1 | 0.00258 | Either |
| $N_{WAG \rightarrow GG}$ | 1 | 0.00258 | Either |
| $N_{WGG \rightarrow AG}$ | 22 | 0.0568 | Between colors |
| $N_{WGG \rightarrow GG}$ | 150 | 0.388 | Within color |

**Table 2.**
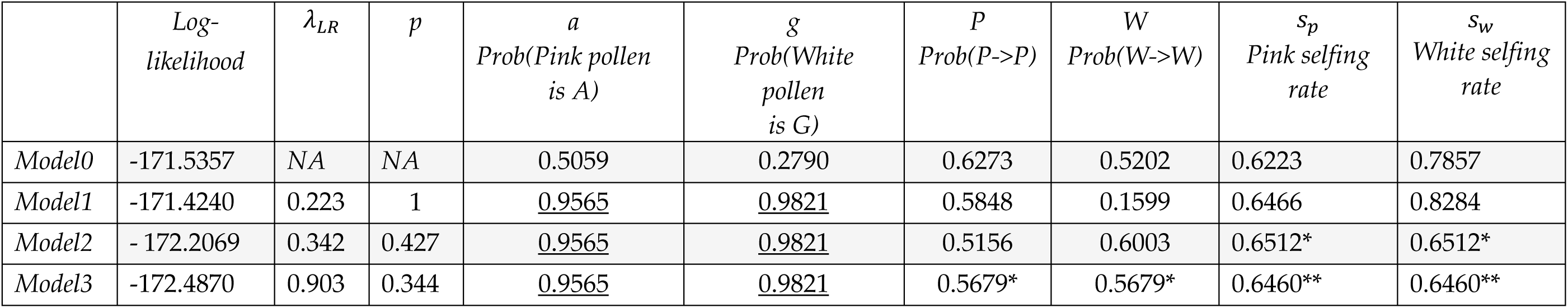
Log-likelihood estimate of siring success. Maximized log-likelihood, likelihood ratio with Model 0 (***λ***_***LR***_), significance (p-value) and parameter estimates (*a, g, P*, *W*, ***s***_***p***_,***s***_***w***_) for the least constrained (Model 0) to most constrained models). Underlined parameters are constrained to a set value, while asterisks indicate parameters are set to be equal. Parameters are defined as follows: *a –* the probability that pink pollen has the A allele; *g -* the probability that white pollen has the G allele; *P* – probability that a pink-limbed plant sires a pink-limbed plant; *W* – probability that a white-limbed plant sires a white-limbed plant; ***s***_***p***_ – selfing rate for pink-limbed plants; ***s***_***w***_ – selfing rate for white-limbed plants

**Table 3.** Color modulates selection on floral traits. ANCOVA summary table for determining the effects of color, corolla length, and sugar concentration (Conc.), and their interaction on seed set and estimated seed number in field RILs. The “Color” effect tests whether there is an overall effect of limb color averaged over all values of flower size or sugar concentration. The “Flower Size” and “Sugar Concentration” main effect indicates whether there is an effect when averaged over both limb colors. The “Flower Size x Color” and “Sugar Concentration X Color” effects test whether the slopes of the regression of fitness on flower size or sugar concentration differ between pink- and white-limbed RILs. Degrees of freedom are 1.

| *Effect* | *Seed Set* |  | *Estimated Seed Number* |  |
| --- | --- | --- | --- | --- |
| | $\chi^2$ | p-value | $\chi^2$ | p-value |
| *Color* | 0.9091 | 0.3404 | 1.3527 | 0.2448 |
| *Flower Size (all RILs)* | 3.0818 | 0.07917 | 1.2327 | 0.2669 |
| *Sugar Concentration (all RILs)* | 3.7763 | 0.05198 | **4.0694** | **0.04367** |
| *Flower Size x Color* | **4.7789** | **0.02881** | **5.6791** | **0.01717** |
| *Sugar Conc x Color* | 3.6246 | 0.056931 | **4.2730** | **0.038722** |

Next, we tested the null hypothesis that selfing rates are equal for pink and white plants (i.e. *s_p_ = s_w_*) by comparing Model 2 (in which both the previous constraint and the constraint of equal selfing rates are included) to Model 1. This comparison also failed to yield a significant decrease in log-likelihood (Table 3), leading us to accept the null hypothesis of equal selfing rates.

Finally, we tested the hypothesis that two probabilities are equal: (1) the probability that a pink plant is sired by a pink plant, and (2) the probability that a white plant is sired by a white plant (i.e. *P* = *W*). We did this by comparing Model 3 (which has the previous two restrictions plus this one) to Model 2. This comparison also fails to yield a significant decrease in log-likelihood (Table 2). This test is equivalent to testing whether per-capita siring success differs between pink and white plants (see Supplementary Methods). Thus, the failure to reject the null hypothesis implies that there was no detectable difference between flower colors in siring success.

### Correlational Selection of Flower Color

Flower color variation modulates the pattern of selection on flower size (corolla length) and sugar concentration in *Ipomoea.* For both traits, there is a positive relationship between the trait and seed mass for pink-limbed plants (p =0.02152 and 0.02036 for corolla length and sugar concentration, respectively), while there is no such relationship for white-limbed plants (Figure 4). This difference in slope is marginally significant for sugar concentration (p =0.056931) and statistically significant for flower size (p =0.02881; Figure 4; Table 3). Similar results are obtained by using seed number instead of total seed mass: both traits are positively correlated with estimated seed number in pink-limbed lines (sugar concentration: p=0.005005; flower size: p=0.07619), but not in white-limbed lines (Figure 3), and this difference in slope is significant for both traits (sugar concentration: p =0.038722; flower size: p=0.01717). Consequently, for at least one of the traits, and likely both, the pattern of selection differs between white and pink RILs.

**Figure 4.**
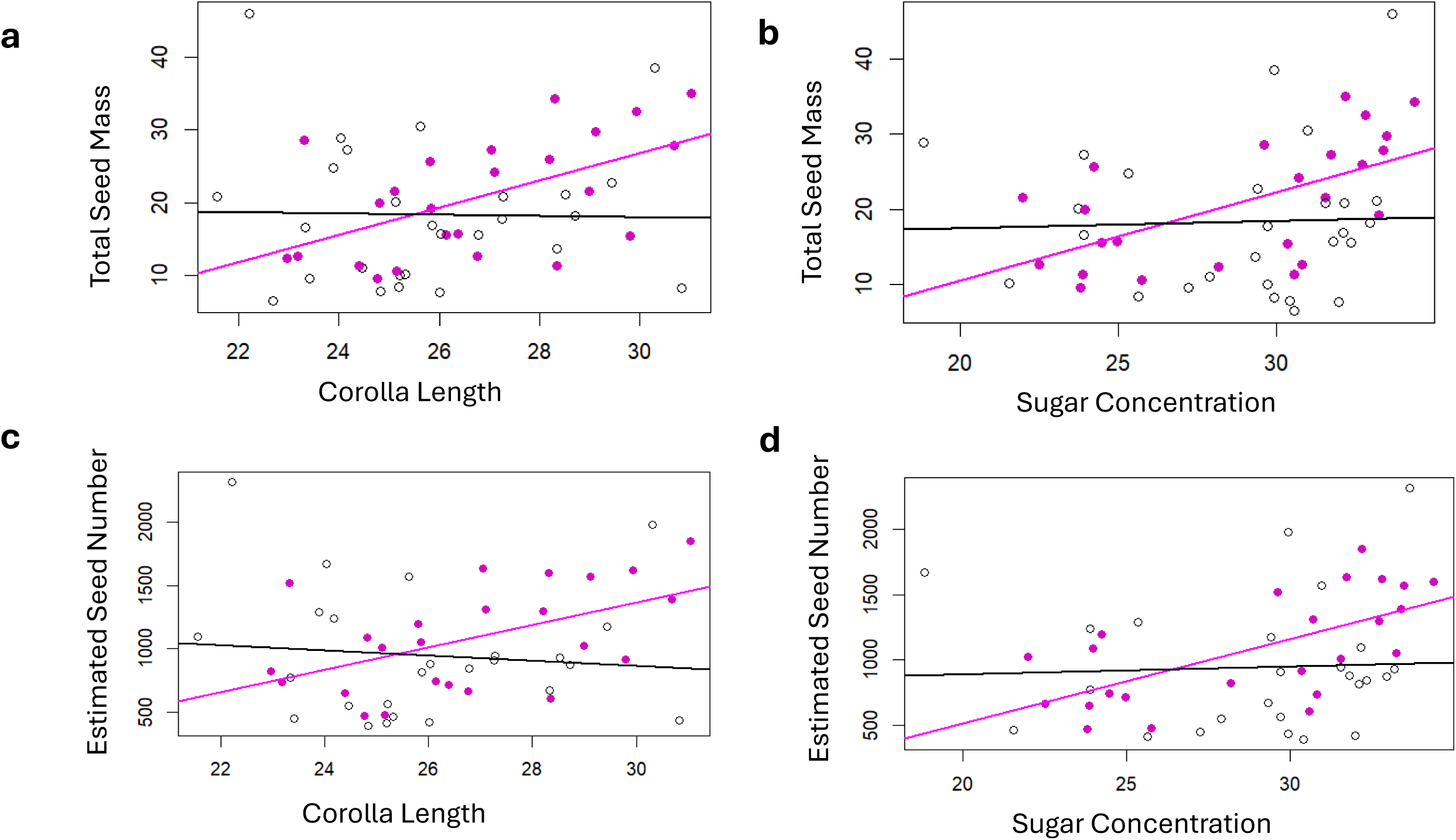
Color Modulates Selection on Floral Traits. The effect of corolla length (left) and sugar concentration (right) on seed set (top) and estimated seed number (bottom) in pink-limbed (pink dots) and white-limbed (white dots) lines. **a-b**). Points are RIL averages for Pink RILs (pink points) and White RILs (open points). For seed mass, linear regressions are significant for pink-limbed (size: 1.878*x-29.508, R^2^ = 0.2736, p-value=0.00513; sugar: 1.177*x-13.069, R^2^ =0.3272, p-value= 0.00207), but not white-limbed lines (size: −0.08003*x+20.39808, R^2^ = −0.03959, p-value=0.922; sugar: 0.09607*x+15.55999, R^2^ = −0.0385, p-value=0.851). **c-d**) Regressions for estimated seed number are significant for flower size and sugar concentration in pink- (size: 88.79*x-1298.26, R^2^ =0.232, p-value=0.00999; sugar: 64.29*x-769.95, R^2^ =0.3871, p= 0.000697) but not white-limbed lines (size: −20.63*x+1481.55, R^2^ =-0.03331, p=0.6387; sugar: 6.041*x+771.144, R^2^ =-0.04128, p=0.8277). The slopes of the lines are equal to the selection gradients for pink- and white-limbed plants multiplied by the genetic variance (V_A_), assuming this variance in these traits is additive.

One possible explanation for these results is that artifactual correlations between flower color, flower size, and sugar concentration had arisen when RILs were created. In this case, lines with larger flowers or sweeter nectar would be more likely to be pink than white, and direct selection on either trait could indirectly impose selection on flower color. However, among RILs, flower color is not detectably associated with either corolla length (F=2.223, DF=49, p-value=0.12) or sugar concentration (F-stat= 0.004696, DF=49, p-value=0.95, Figures 4, S9). Moreover, flower size is only loosely correlated with sugar concentration (F-stat=6.04, DF=47, p-value=0.018, Adj R-squared=0.0916, Figure S10); these numbers are consistent with the low genetic correlations among a larger subset of lines between nectary sugar and flower size traits (LIAO *et al*. 2022).

### Correlational selection on flower color due to modulation of selection on corolla length and sugar concentration

These patterns demonstrate the operation of epistatic selection: the pattern of selection operating on each of the floral traits depends on flower color. Here we employ evolutionary modelling to infer whether this epistatic selection could cause purifying selection against the color allele. Using flower size as the quantitative trait (see Methods for model parameter values) and total seed mass as the fitness measure, we ran our genetic model using different values of migration rate, *m*, between 0.002 and 0.04 to determine the relationship between migration rate and equilibrium color locus allele frequencies.

If the introgression rate is less than a critical value (between 0.032 and 0.034), the frequency of the “pink” allele reaches an equilibrium that decreases as *m* increases (Figure **5a**), and mean phenotype reaches an equilibrium less than the maximum possible (Figure **5b**, Figure S11). With *m* ≤ 0.01, introgression is effectively prevented. Conversely, when *m* is greater than the critical value (∼0.032), repeated introgression fixes the “white” allele (Figure **5a**) and reduces the frequency of any quantitative trait (i.e., size allele) to its minimum (Figure S11). We find that these patterns are qualitatively consistent across parameter values representing all four panels of Figure 4 (see Figures S12-14).

**Figure 5.**
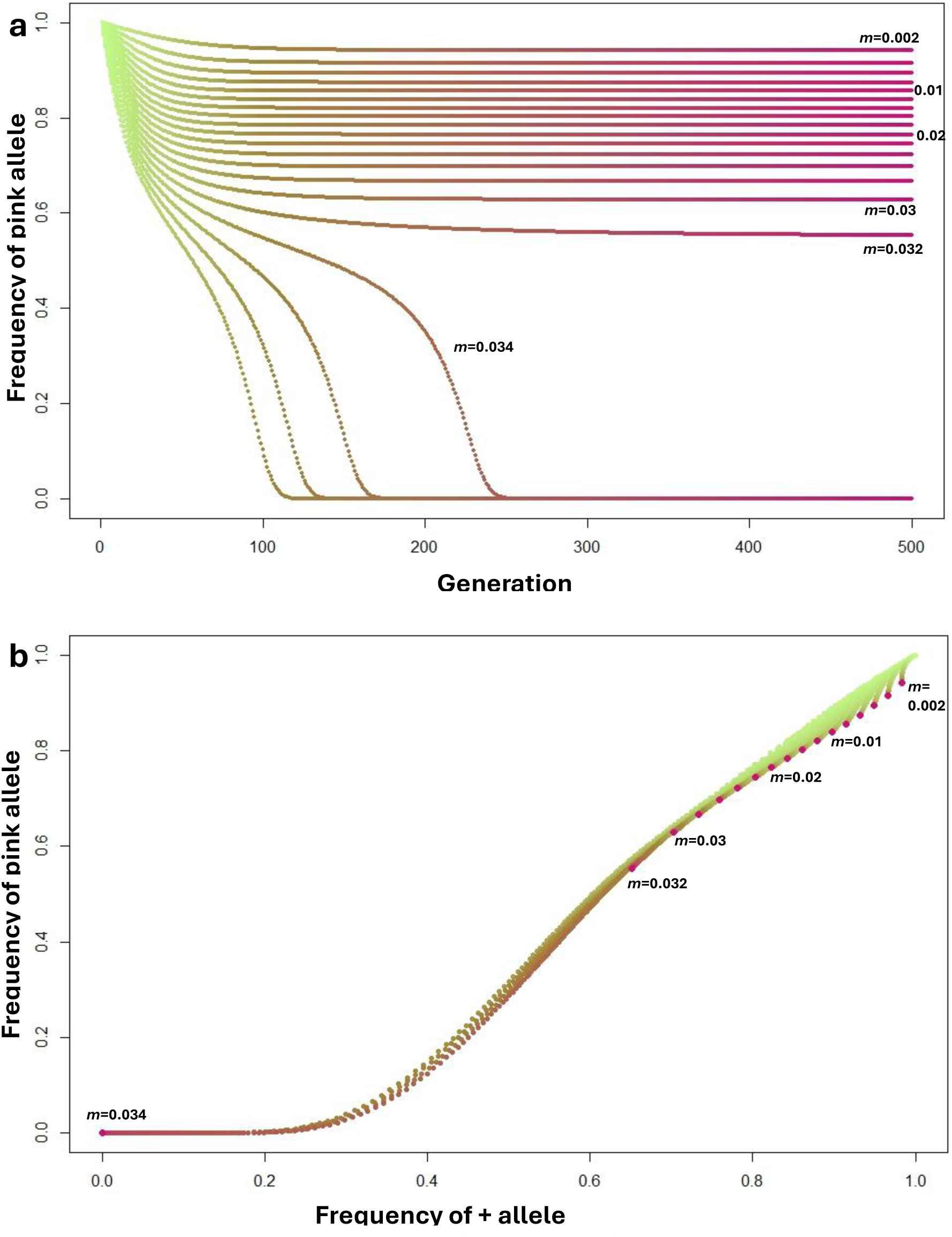
Joint evolution of flower color and quantitative trait from explicitly genetic model. **a)** The change in the frequency of the “pink” allele over 500 generations given continual gene flow from *I. lacunosa* into *I. cordatotriloba*. Migration rate (*m*) corresponding to the top trajectory is 0.002 and increases by 0.002 for each subsequent lower trajectory. **b)** Trajectory of “pink” allele and + (*I.* cordata) size alleles given different rates of migration. Note that because all quantitative trait loci are equivalent, the trajectory is the same at all loci. Trajectories start with fixation of *I. cordatotriloba* alleles and end at red points. Migration rates as in part A. All migration rates greater than 0.032 result in fixation of the introgressing (*I. lacunosa*) haplotype, while lower values allow color and size alleles to resist introgression via correlational selection.

We additionally asked if this epistatic selection between color and trait loci can lead to linkage disequilibrium (LD) and thus strengthen genetic correlations among color and quantitative trait loci. We find that the LD generated is quite small and increases the genetic correlation among the traits by only at around 3% of the maximum correlation possible (r_G_ =0.02944). This suggests that most of the selection favoring an increase in the pink allele when the mean trait value is high is directly due to epistatic selection, rather than due to indirect selection through a genetic correlation between the trait and color. To quantify the proportions of allele frequency change at the color locus that are due to correlational selection vs. indirect selection due to LD, we calculated the allele frequency changes due to selection at equilibrium as done before but artificially removed LD and observed how the allele frequency changes due to selection (see Methods). Using model parameters from Figure 4D, and a migration rate value of *m*=0.04, the frequency of the pink allele changes by 0.273 when LD is present, and by 0.0231 when LD is removed. As a result, we find that 84.6% of the change in allele frequency due to selection is due to correlational selection and 15.4% is due to indirect section from generated LD. So, while the pink allele resists introgression via correlational selection, the increase in LD from epistatic selection can amplify the overall response to selection at the color locus.

## Discussion

The objective of this study was to determine whether purifying selection acts to prevent introgression at the flower-color locus from *I. lacunosa* into *I. cordatotriloba*. We first investigated whether selection acts directly against the white allele in an experimental population of *I. cordatotriloba* by asking if pink-limbed lines outperformed white-limbed lines across any component of lifetime fitness – germination, survival, fecundity, or siring success. We found no evidence of such selection for any of these four fitness components on floral color. This result contrasts with previous studies that have found evidence for direct selection at loci corresponding to key traits that likely prevent subsequent introgression (Bradshaw et al. 1998, Hodges et al. 2002, Hopkins and Rausher 2012, Kenney and Sweigart 2016, Farnitano et al. 2025).

Notably, most of these studies attempted to quantify only direct selection at the loci examined. While direct selection may play a role in preventing introgression, it does not preclude other forms of selection, such as correlational selection, from generating resistance to introgression. Our research specifically demonstrates one such case of correlational selection – in *Ipomoea* the genotype of one trait (floral color) modulates the strength and direction of selection on quantitative traits (flower size and sugar concentration). Because the fitness of pink-limbed plants increases with the value of the quantitative trait while that of white-limbed plants does not, the fitness of pink-limbed individuals will be higher than that of white-limbed individual with the same trait value when this trait value is greater than a critical value. As long as the trait mean is above this threshold, as will be the case in sympatric *I. cordatotriloba* populations, this fitness difference favors an increase in the frequency of the pink allele.

Our model indicates that when gene flow from *I. lacunosa* is below a critical level, a polymorphic equilibrium will be reached at the color locus. This equilibrium reflects a balance between correlational selection increasing the pink allele frequency and migration decreasing it. Notably, the model indicates that the trait that modulates selection need not be directly selected for introgression to be prevented. Our results are consistent with the finding that correlational selection, in conjunction with direct selection, prevents hybridization between *Ipomopsis* species that differ in flower color, and thus acts to limit flower-color introgression (Meléndez-Ackerman et al. 1997). The combination of these results and our findings further support prior calls (see Campbell 2009, Svensson et al. 2021) to integrate research on correlational selection with that on mating system evolution.

Our results do not provide a clear mechanism as to why a shift from white to pink flower color induces directional selection for other floral traits. One possible explanation is that pollinators visit plants in a way that mirrors fitness results, with visits correlating with flower size and nectar sugar concentration when plants are pink- but not white-limbed. However, a separate experiment found no evidence supporting this hypothesis (Colen 2025). The mechanisms underlying this epistatic selection are therefore unclear, and further field experiments are needed to test the extent to which pollinators, pleiotropic effects, herbivory, or heat stress can generate observed patterns of selection.

Our model indicates that while correlational selection can limit introgression if gene flow is less than a critical value and that the equilibrium frequency of the introgressing allele increases as migration rate increases. Above that critical value, gene flow overpowers selection and leads to complete introgression. A previous survey of 6 sympatric populations yielded an average frequency the white allele in sympatric populations of *I. cordatotriloba* of 0.078 (Duncan 2013); based on our model, this corresponds to an average migration rate of approximately 0.002. A migration rate this low seems reasonable given the strong cross incompatibilities between *I. cordatotriloba* and *I. lacunosa* (Ostevik et al. 2021, Rifkin et al. 2023).

Given the estimated migration rate at which the color locus can resist introgression, we believe that these results are likely generalizable across hybridizing species. In many cases, between-species gene flow may be weak enough to allow correlational selection to prevent introgression; approximately 2/3 of sympatric species pairs exhibit either strong prezygotic isolation (>75% displacement in reproductive overlap), strong postzygotic isolation (>75% fitness advantage of parental species as compared to hybrid offspring), or both (Christie et al. 2022). This suggests that one of the conditions required for correlational selection to prevent introgression is present in many systems, and implies that, when genetic migration is low, the state of one allele can play a substantial role in determining whether many other loci simultaneously resist introgression simultaneously.

Although we failed to find evidence of direct purifying selection opposing introgression of the white allele in sympatric *I. cordatotriloba*, several caveats prevent us from definitively ruling out direct selection on flower color. First, while we find no effect of color on germination or early growth, the spacing between plants prevented us from explicitly assessing if the flower color allele imparts a competitive advantage in accessing limited early-season resources such as light or water. Because flower color often confers a fitness advantage in drought conditions (Warren and Mackenzie 2001, Schemske and Bierzychudek 2007, Vaidya et al. 2018), increased competition for limited water resources could lead to a fitness advantage for the pink allele. Second, because plants were drip irrigated during transplanting, we cannot determine if flower color affects drought tolerance during early growth. Third, flower color may provide a fitness advantage in anomalous years when abiotic (rainfall, temperature) or biotic (herbivory, pollinator abundance) conditions not seen during our field season occur. Fourth, we were unable to assess if offspring of white-limbed plants were more likely to be sired by *I. lacunosa* than offspring of pink-limbed plants, leading to a potential fitness advantage for pink-limbed plants by reducing costly interspecific pollination. However, even if pink color is directly selected via one of these components, our results demonstrate that color’s resistance to introgression is at least in part explainable by correlational selection.

In summary, our results show that correlational selection involving a binary and quantitative trait can contribute to resistance to introgression at the binary-trait locus. As a corollary, these results suggest that the extent that quantitative alleles resist introgression depends upon the state of the binary trait locus. Correlational selection of this type thus provides a novel mechanism to explain how introgression across multiple sites may simultaneously resist introgression via modulation by key binary-trait loci. While interspecific incompatibilities (Rieseberg et al. 1999, Hohenlohe et al. 2012, Schumer et al. 2014, Pool 2015), and assortative mating (Bradshaw Jr et al. 1995, Hopkins and Rausher 2012, Fogel 2022, Schield et al. 2024) have clearly shaped the genomic landscape of loci that resist introgression, correlational selection of the type we describe provides a novel mechanism to “package” together loci across the genome that resist introgression. Additionally, our results emphasize that correlated responses to selection need not be due solely to genetic correlations between non-incompatibility and incompatibility loci (Juenger et al. 2005, Conner et al. 2011, Wessinger et al. 2014, Smith 2016), and can be generated specifically by the mode of selection. We suspect that this type of correlational selection is more common than previously reported. Further investigation on the role of correlated selection on introgression will allow us to better understand how species can remain distinct despite hybridization.

## Acknowledgements

We would like to thank Tanya Duncan, Joanna Rifkin, and Irene Liao whose prior research on *Ipomoea cordatotriloba* and *lacunosa* made this project possible. We would like to especially thank Irene Liao for the extensive data collected on floral and nectary traits that were instrumental to all analyses conducted on correlational selection. We would also like to thank Drs. John Willis, William Morris, Karin Pfennig, and Craig Lowe for feedback on research analyses and on earlier versions of this manuscript. Field research would not have been possible without the assistance provided by staff at the Border Belt Tobacco Research Station, particularly superintendent Lloyd Ransom. We would like to thank undergraduates Katie Meehl, Lola Adewale, Porter Petruzziello, and Caroline Metz for their assistance with data collection. Funding for this project was generously provided via the following sources: North Carolina Native Plant Society, Sigma Xi, and the Duke Biology Department.

## Author contributions

J.Z.C. collected all field data on germination rate, survival, and fecundity and conducted analyses of parentage for assessing siring rate. J.Z.C. provided code for mathematical models built in R while M.D.R. provided code for those built in Mathematica. J.Z.C. and M.D.R. conceived of the projected, designed the experiment, performed analyses, and wrote the paper.

## Data availability

Data used for analyses will be made available upon a digital repository (Dryad) upon publication. All code will be made available on github upon acceptance of the manuscript for publication.

## References

Barton, N. H., and M. A. R. De Cara. 2009. THE EVOLUTION OF STRONG REPRODUCTIVE ISOLATION. Evolution 63:1171–1190.

Bates, D., M. Mächler, B. Bolker, and S. Walker. 2015. Fitting linear mixed-effects models using lme4, v1.1.36. Journal of statistical software 67:1–48.

Bradshaw, H., K. G. Otto, B. E. Frewen, J. K. McKay, and D. W. Schemske. 1998. Quantitative trait loci affecting differences in floral morphology between two species of monkeyflower (Mimulus). Genetics 149:367–382.

Bradshaw Jr, H., S. M. Wilbert, K. Otto, and D. Schemske. 1995. Genetic mapping of floral traits associated with reproductive isolation in monkeyflowers (Mimulus). Nature 376:762–765.

Campbell, D. R. 2009. Using phenotypic manipulations to study multivariate selection of floral trait associations. Annals of Botany 103:1557–1566.

Carruthers, T., P. Muñoz-Rodríguez, J. R. I. Wood, and R. W. Scotland. 2020. The temporal dynamics of evolutionary diversification in Ipomoea. Molecular Phylogenetics and Evolution 146:106768.

Christie, K., L. S. Fraser, and D. B. Lowry. 2022. The strength of reproductive isolating barriers in seed plants: Insights from studies quantifying premating and postmating reproductive barriers over the past 15 years. Evolution 76:2228–2243.

Clements, F. E., and G. W. Goldsmith. 1924. The phytometer method in ecology: the plant and community as instruments. Carnegie Institution of Washington.

Colen, J. Z. 2025. Investigating How the Color Allele in Ipomoea cordatotriloba Resists Introgression via Epistatic Selection. Ph.D. Duke University, United States -- North Carolina.

Comeault, A. A., S. M. Flaxman, R. Riesch, E. Curran, V. Soria-Carrasco, Z. Gompert, T. E. Farkas, M. Muschick, T. L. Parchman, and T. Schwander. 2015. Selection on a genetic polymorphism counteracts ecological speciation in a stick insect. Current Biology 25:1975–1981.

Conner, J. K., K. Karoly, C. Stewart, V. A. Koelling, H. F. Sahli, and F. H. Shaw. 2011. Rapid independent trait evolution despite a strong pleiotropic genetic correlation. The American Naturalist 178:429–441.

Conner, J. K., and S. Rush. 1996. Effects of flower size and number on pollinator visitation to wild radish, Raphanus raphanistrum. Oecologia 105:509–516.

Darwin, C. 1859. On the origin of species by means of natural selection, or preservation of favoured races in the struggle for life. London: John Murray, 1859.

Delgado, T., L. C. Leal, J. H. L. El Ottra, V. L. G. Brito, and A. Nogueira. 2023. Flower size affects bee species visitation pattern on flowers with poricidal anthers across pollination studies. Flora 299:152198.

Dietrich, A. L., C. Nilsson, and R. Jansson. 2013. Phytometers are underutilised for evaluating ecological restoration. Basic and Applied Ecology 14:369–377.

Duncan, T. M. 2013. The Mating System Evolution of Ipomoea lacunosa. Citeseer.

Duncan, T. M., and M. D. Rausher. 2013. Evolution of the selfing syndrome in Ipomoea. Frontiers in Plant Science **Volume** 4 - 2013.

Duncan, T. M., and M. D. Rausher. 2020. Selection favors loss of floral pigmentation in a highly selfing morning glory. PLoS One 15:e0231263.

Farnitano, M. C., K. Karoly, and A. L. Sweigart. 2025. Fluctuating reproductive isolation and stable ancestry structure in a fine-scaled mosaic of hybridizing Mimulus monkeyflowers. PLoS genetics 21:e1011624.

Fehr, C., and M. D. Rausher. 2004. Effects of variation at the flower-colour A locus on mating system parameters in Ipomoea purpurea. Molecular ecology 13:1839–1847.

Felsenstein, J. 1981. Skepticism Towards Santa Rosalia, or Why are There so Few Kinds of Animals? Evolution 35:124–138.

Finley, A., S. Banerjee, Ø. Hjelle, and R. Bivand. 2024. MBA: multilevel B-spline approximation. R package version 0.1–2.

Fishman, L., A. L. Sweigart, A. M. Kenney, and S. Campbell. 2014. Major quantitative trait loci control divergence in critical photoperiod for flowering between selfing and outcrossing species of monkeyflower (M imulus). New Phytologist 201:1498–1507.

Fogel, A. S. 2022. Genomic and Phenotypic Consequences of Hybridization in Wild Baboons. Ph.D. Duke University, United States -- North Carolina.

Fornoff, F., A. M. Klein, F. Hartig, G. Benadi, C. Venjakob, H. M. Schaefer, and A. Ebeling. 2017. Functional flower traits and their diversity drive pollinator visitation. Oikos 126:1020–1030.

Fowler, R. E., E. L. Rotheray, and D. Goulson. 2016. Floral abundance and resource quality influence pollinator choice. Insect Conservation and Diversity 9:481–494.

Fox, J., and S. Weisberg. 2018. An R companion to applied regression; version 3.1.3. Sage publications.

Friedman, J., A. D. Twyford, J. H. Willis, and B. K. Blackman. 2015. The extent and genetic basis of phenotypic divergence in life history traits in Mimulus guttatus. Molecular ecology 24:111–122.

Fulton, M., and S. A. Hodges. 1999. Floral isolation between Aquilegia formosa and Aquilegia pubescens. Proceedings of the Royal Society of London. Series B: Biological Sciences 266:2247–2252.

Gavrilets, S. 1997. HYBRID ZONES WITH DOBZHANSKY-TYPE EPISTATIC SELECTION. Evolution 51:1027–1035.

Gavrilets, S., and A. Hastings. 1996. Founder Effect Speciation: A Theoretical Reassessment. The American Naturalist 147:466–491.

Hankin, C. M. 2024. Genomic Insights Into Species Divergence: Unraveling Genetic Architecture Amid Gene Flow in the Mimulus guttatus Complex. Tulane University.

Hawthorne, D. J., and S. Via. 2001. Genetic linkage of ecological specialization and reproductive isolation in pea aphids. Nature 412:904–907.

Hodges, S. A., J. B. Whittall, M. Fulton, and J. Y. Yang. 2002. Genetics of Floral Traits Influencing Reproductive Isolation between Aquilegia formosa and Aquilegia pubescens. . The American Naturalist 159:S51–S60.

Hohenlohe, P. A., S. Bassham, M. Currey, and W. A. Cresko. 2012. Extensive linkage disequilibrium and parallel adaptive divergence across threespine stickleback genomes. Philosophical Transactions of the Royal Society B: Biological Sciences 367:395–408.

Hopkins, R., and M. D. Rausher. 2012. Pollinator-mediated selection on flower color allele drives reinforcement. Science 335:1090–1092.

Juenger, T., J. M. Pérez-Pérez, S. Bernal, and J. L. Micol. 2005. Quantitative trait loci mapping of floral and leaf morphology traits in Arabidopsis thaliana: evidence for modular genetic architecture. Evolution & development 7:259–271.

Kenney, A. M., and A. L. Sweigart. 2016. Reproductive isolation and introgression between sympatric Mimulus species. Molecular ecology 25:2499–2517.

Lande, R., and S. J. Arnold. 1983. The measurement of selection on correlated characters. Evolution:1210–1226.

Langsrud, Ø. 2003. ANOVA for unbalanced data: Use Type II instead of Type III sums of squares. Statistics and computing 13:163–167.

Liao, I. T., J. L. Rifkin, G. Cao, and M. D. Rausher. 2022. Modularity and selection of nectar traits in the evolution of the selfing syndrome in Ipomoea lacunosa (Convolvulaceae). New Phytologist 233:1505–1519.

Lindtke, D., and C. A. Buerkle. 2015. The genetic architecture of hybrid incompatibilities and their effect on barriers to introgression in secondary contact. Evolution 69:1987–2004.

Mallet, J. 2005. Hybridization as an invasion of the genome. Trends in ecology & evolution 20:229–237.

Mallinger, R., and J. Prasifka. 2017. Bee visitation rates to cultivated sunflowers increase with the amount and accessibility of nectar sugars. Journal of Applied Entomology 141:561–573.

Meléndez-Ackerman, E., D. R. Campbell, and N. M. Waser. 1997. Hummingbird behavior and mechanisms of selection on flower color in Ipomopsis. Ecology 78:2532–2541.

Meléndez-Ackerman, E., and D. R. Campbell. 1998. Adaptive significance of flower color and inter-trait correlations in an Ipomopsis hybrid zone. Evolution 52:1293–1303.

Murray, M., and W. Thompson. 1980. Rapid isolation of high molecular weight plant DNA. Nucleic acids research 8:4321–4326.

Nottebrock, H., B. Schmid, K. Mayer, C. Devaux, K. J. Esler, K. Böhning-Gaese, M. Schleuning, J. Pagel, and F. M. Schurr. 2017. Sugar landscapes and pollinator-mediated interactions in plant communities. Ecography 40:1129–1138.

Ostevik, K. L., J. L. Rifkin, H. Xia, and M. D. Rausher. 2021. Morning glory species co-occurrence is associated with asymmetrically decreased and cascading reproductive isolation. Evolution Letters 5:75–85.

Payseur, B. A. 2010. Using differential introgression in hybrid zones to identify genomic regions involved in speciation. Molecular Ecology Resources 10:806–820.

Payseur, B. A., and L. H. Rieseberg. 2016. A genomic perspective on hybridization and speciation. Molecular Ecology 25:2337–2360.

Pool, J. E. 2015. The mosaic ancestry of the Drosophila genetic reference panel and the D. melanogaster reference genome reveals a network of epistatic fitness interactions. Molecular biology and evolution 32:3236–3251.

R Core Team. 2024. R: A Language and Environment for Statistical Computing. R Foundation for Statistical Computing, Vienna, Austria.

Rausher, M. D., D. Augustine, and A. VanderKooi. 1993. ABSENCE OF POLLEN DISCOUNTING IN A GENOTYPE OF IPOMOEA PURPUREA EXHIBITING INCREASED SELFING. Evolution 47:1688–1695.

Rausher, M. D., and J. D. Fry. 1993. Effects of a locus affecting floral pigmentation in Ipomoea purpurea on female fitness components. Genetics 134:1237–1247.

Rieseberg, L. H., M. A. Archer, and R. K. Wayne. 1999. Transgressive segregation, adaptation and speciation. Heredity 83:363–372.

Rieseberg, L. H., and J. F. Wendel. 1993. Introgression and its consequences.

Rifkin, J. L., G. Cao, and M. D. Rausher. 2021. Genetic architecture of divergence: the selfing syndrome in Ipomoea lacunosa. American Journal of Botany 108:2038–2054.

Rifkin, J. L., A. S. Castillo, I. T. Liao, and M. D. Rausher. 2019a. Gene flow, divergent selection and resistance to introgression in two species of morning glories (Ipomoea). Molecular ecology 28:1709–1729.

Rifkin, J. L., I. T. Liao, A. S. Castillo, and M. D. Rausher. 2019b. Multiple aspects of the selfing syndrome of the morning glory Ipomoea lacunosa evolved in response to selection: A Qst-Fst comparison. Ecology and Evolution 9:7712–7725.

Rifkin, J. L., K. L. Ostevik, and M. D. Rausher. 2023. Complex cross-incompatibility in morning glories is consistent with a role for mating system in plant speciation. Evolution 77:1691–1703.

Schemske, D. W., and P. Bierzychudek. 2007. SPATIAL DIFFERENTIATION FOR FLOWER COLOR IN THE DESERT ANNUAL LINANTHUS PARRYAE: WAS WRIGHT RIGHT? Evolution 61:2528–2543.

Schield, D. R., J. K. Carter, E. S. C. Scordato, I. I. Levin, M. R. Wilkins, S. A. Mueller, Z. Gompert, P. Nosil, J. B. W. Wolf, and R. J. Safran. 2024. Sexual selection promotes reproductive isolation in barn swallows. Science 386:eadj8766.

Schumer, M., R. Cui, D. L. Powell, R. Dresner, G. G. Rosenthal, and P. Andolfatto. 2014. High-resolution mapping reveals hundreds of genetic incompatibilities in hybridizing fish species. Elife 3:e02535.

Seehausen, O., R. K. Butlin, I. Keller, C. E. Wagner, J. W. Boughman, P. A. Hohenlohe, C. L. Peichel, G.-P. Saetre, C. Bank, Å. Brännström, A. Brelsford, C. S. Clarkson, F. Eroukhmanoff, J. L. Feder, M. C. Fischer, A. D. Foote, P. Franchini, C. D. Jiggins, F. C. Jones, A. K. Lindholm, K. Lucek, M. E. Maan, D. A. Marques, S. H. Martin, B. Matthews, J. I. Meier, M. Möst, M. W. Nachman, E. Nonaka, D. J. Rennison, J. Schwarzer, E. T. Watson, A. M. Westram, and A. Widmer. 2014. Genomics and the origin of species. Nature Reviews Genetics 15:176–192.

Slatkin, M. 2008. Linkage disequilibrium — understanding the evolutionary past and mapping the medical future. Nature Reviews Genetics 9:477–485.

Smith, S. D. 2016. Pleiotropy and the evolution of floral integration. New Phytologist 209:80–85.

Soetaert, K., and M. K. Soetaert. 2024. Package ‘plot3D’. Tools for plotting 3.

Stephenson IV, D. O., L. R. Oliver, N. R. Burgos, and E. E. Gbur Jr. 2006. Identification and characterization of pitted morningglory (Ipomoea lacunose) ecotypes. Weed Science 54:78–86.

Suarez-Gonzalez, A., C. Lexer, and Q. C. Cronk. 2018. Adaptive introgression: a plant perspective. Biology letters 14:20170688.

Svensson, E. I., S. J. Arnold, R. Bürger, K. Csilléry, J. Draghi, J. M. Henshaw, A. G. Jones, S. De Lisle, D. A. Marques, and K. McGuigan. 2021. Correlational selection in the age of genomics. Nature ecology & evolution 5:562–573.

Taylor, S. A., and E. L. Larson. 2019. Insights from genomes into the evolutionary importance and prevalence of hybridization in nature. Nature ecology & evolution 3:170–177.

Vaidya, P., A. McDurmon, E. Mattoon, M. Keefe, L. Carley, C.-R. Lee, R. Bingham, and J. T. Anderson. 2018. Ecological causes and consequences of flower color polymorphism in a self-pollinating plant (Boechera stricta). New Phytologist 218:380–392.

Warren, J., and S. Mackenzie. 2001. Why are all colour combinations not equally represented as flower-colour polymorphisms? New Phytologist 151:237–241.

Wessinger, C. A., L. C. Hileman, and M. D. Rausher. 2014. Identification of major quantitative trait loci underlying floral pollination syndrome divergence in Penstemon. Philosophical Transactions of the Royal Society B: Biological Sciences 369:20130349.

Wheat, C. W., H. W. Fescemyer, J. Kvist, E. Tas, J. C. Vera, M. J. Frilander, I. Hanski, and J. H. Marden. 2011. Functional genomics of life history variation in a butterfly metapopulation. Molecular ecology 20:1813–1828.

Wu, C.-I. 2001. The genic view of the process of speciation. Journal of Evolutionary Biology 14:851–865.

