## Supplementary Figures and Tables for "Flower color locus resists introgression due to correlational selection with other floral traits in *Ipomoea cordatotriloba*"

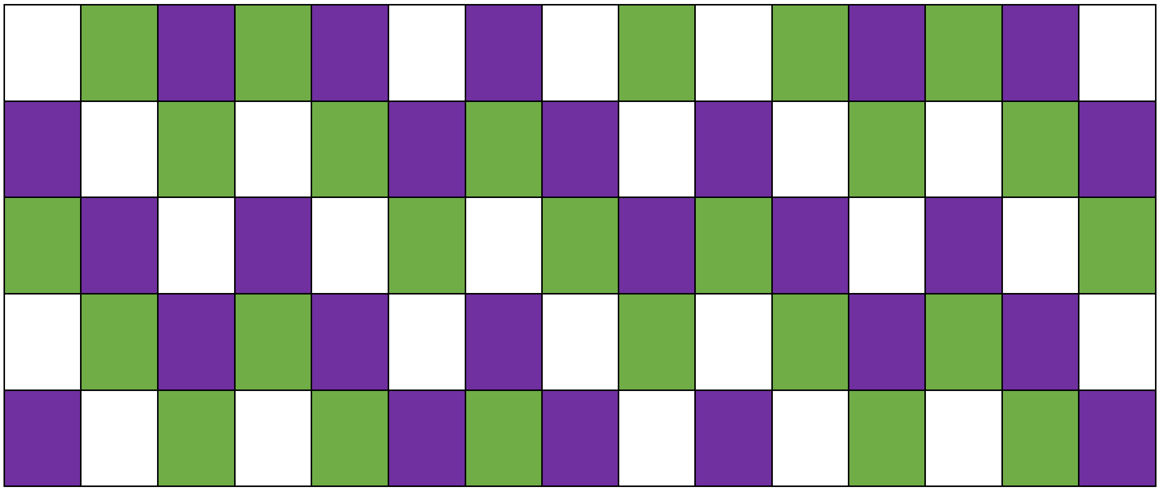

**Figure S1. Field Layout Schematic for a Representative Plot.** Color-coding represents locations for plants of the same type for three types– pink-limbed lines, white-limbed lines, and *I. lacunosa*. Note that the colors are not coordinated across plots (i.e. green boxes may represent pink-limbed lines in one plot, but white-limbed lines or lac in a separate plot). RIL identity is randomized within type for pink-limbed and white-limbed lines.

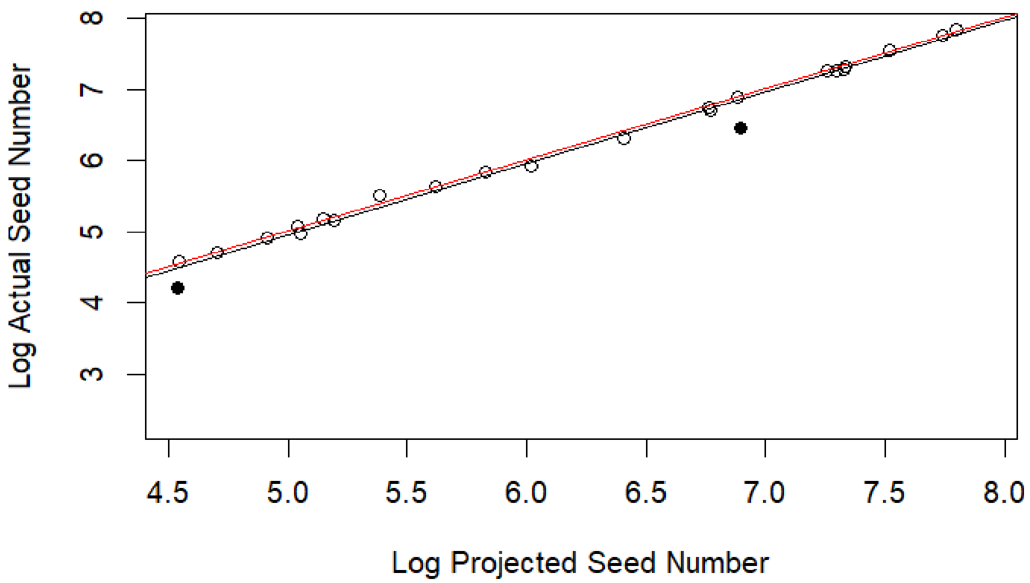


**Figure S2. Estimated seed number highly predicts seed counted** ($p<2*10-16$, $R2=0.9883$). The black trend line tightly corresponds to the red 1/1 trend line. Results are presented as log seed number for clearer visibility. Filled circles represent outliers

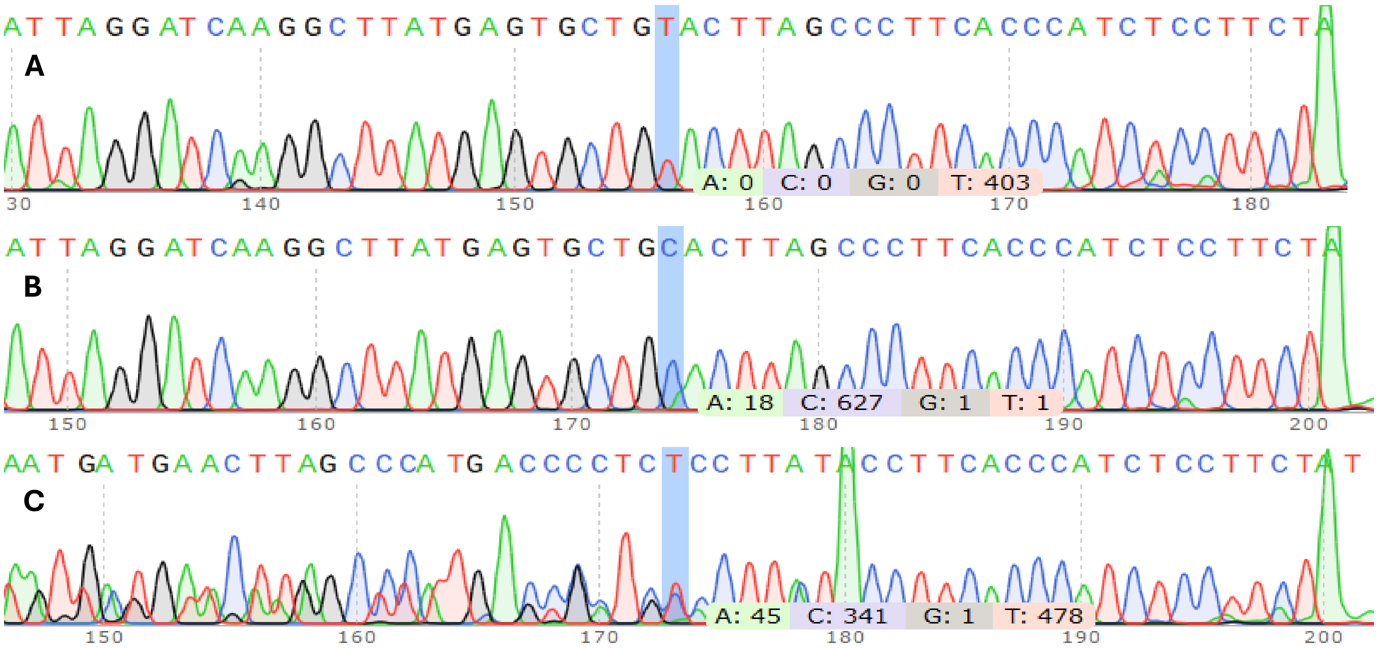


**Figure S3 Example chromatographs for homozygotes of the allele associated with pink-limbed lines (A), white-limbed lines (B), and heterozygotes** (**C**). Blue bar indicates position of the focal SNP used as a marker. Note that homozygotes have distinct frequencies of the T (**A**) and C **(B)** alleles, while heterozygotes have approximately similar readings for both alleles. Additionally, homozygotes of the pink-limbed allele have a peak for nucleotide A around base 180 (**A**), white-limbed allele has a similar peak around base 200 (**B**), and heterozygotes have two distinct peaks, one at each site, along with doubled chromatograph sequence consistent with heterozygous sequence preceding the peak at base 180 (**C**).

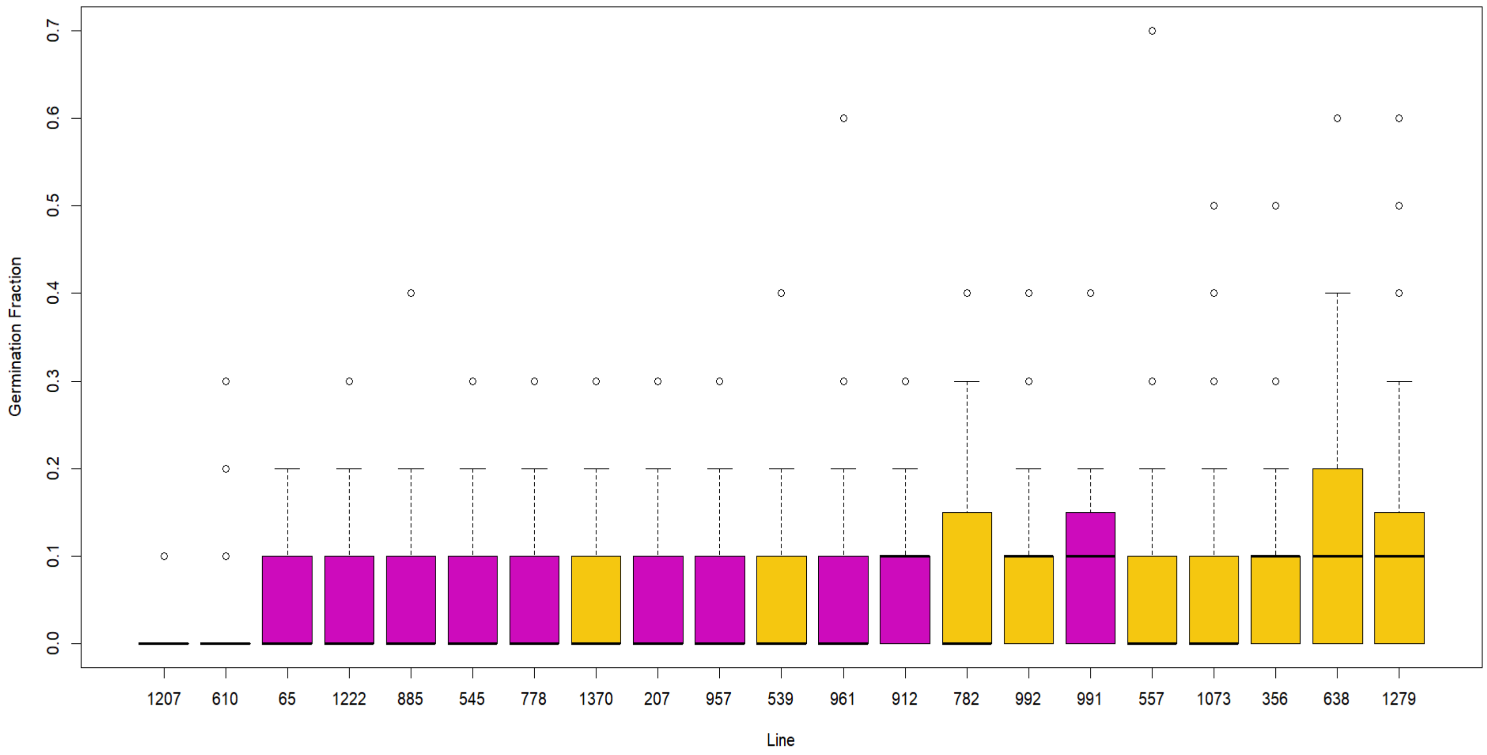


**Figure S4. Germination probability is low for all lines.** Each bar is average proportion of 10 seeds that germinated for each RIL. Pink bars: pink-limbed RILs. Yellow bars: white-limbed RILs. Numbers on x-axis are RIL identification numbers. Black bars represent median germination fraction, whiskers represent 1.5 times the interquartile range, and empty circles represent all outliers beyond this range.

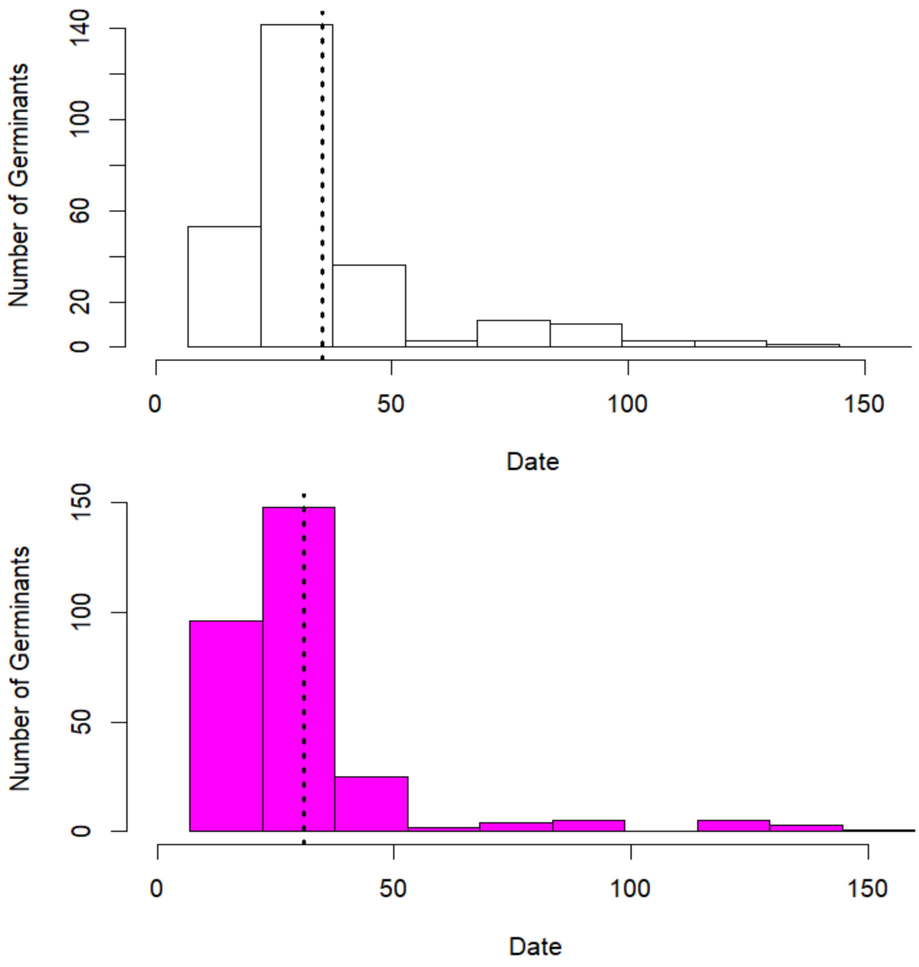


**Figure S5. Pink limbed lines germinate earlier than white-limbed lines.** Histograms of germinant number with time for white-limbed (top) and pink-limbed (bottom, pink) plants. Vertical dashed lines are mean days after planting (Date) for germination of white-limbed (35.45) and pink-limbed (31.03) lines

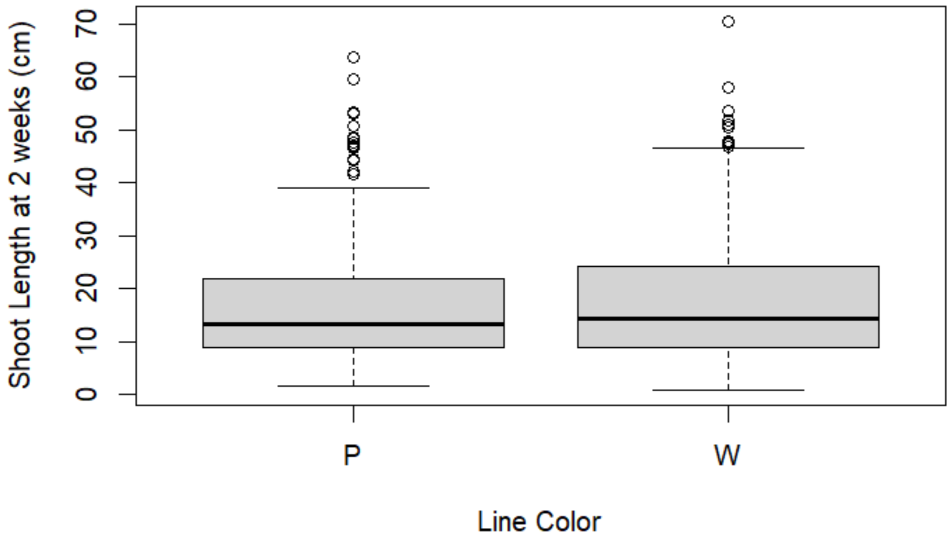


**Figure S6**. **No difference in shoot length 2 weeks after transplant (a proxy for early growth rate) between pink(P) and white-limbed (W) lines.** Black lines represent median shoot length for each color, whiskers represent 1.5 times the interquartile range (grey boxes, middle 50% of data range), and circles represent outliers outside this range.

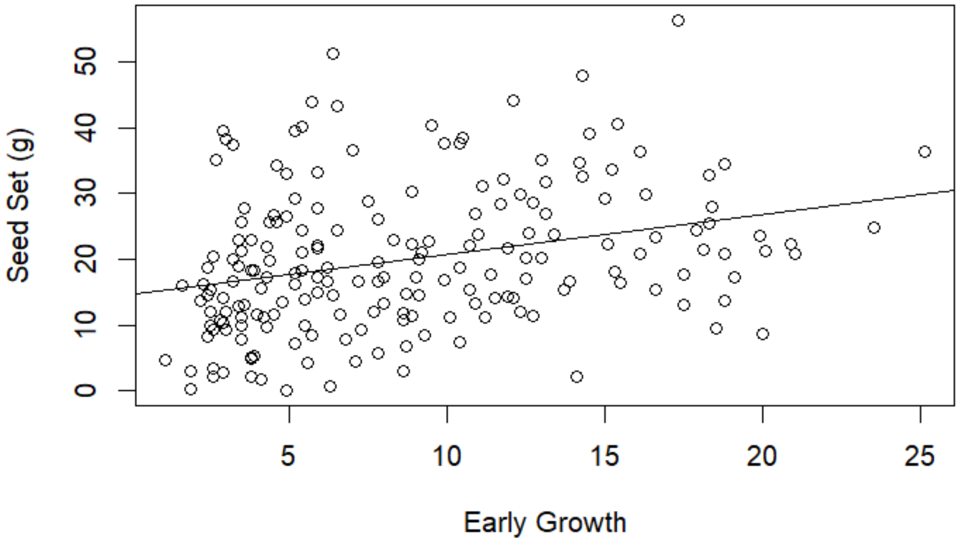


**Figure S7. Early growth is weakly correlated with seed set.** ($p=2.855*10^{-5}$, $R^2=0.08433$).

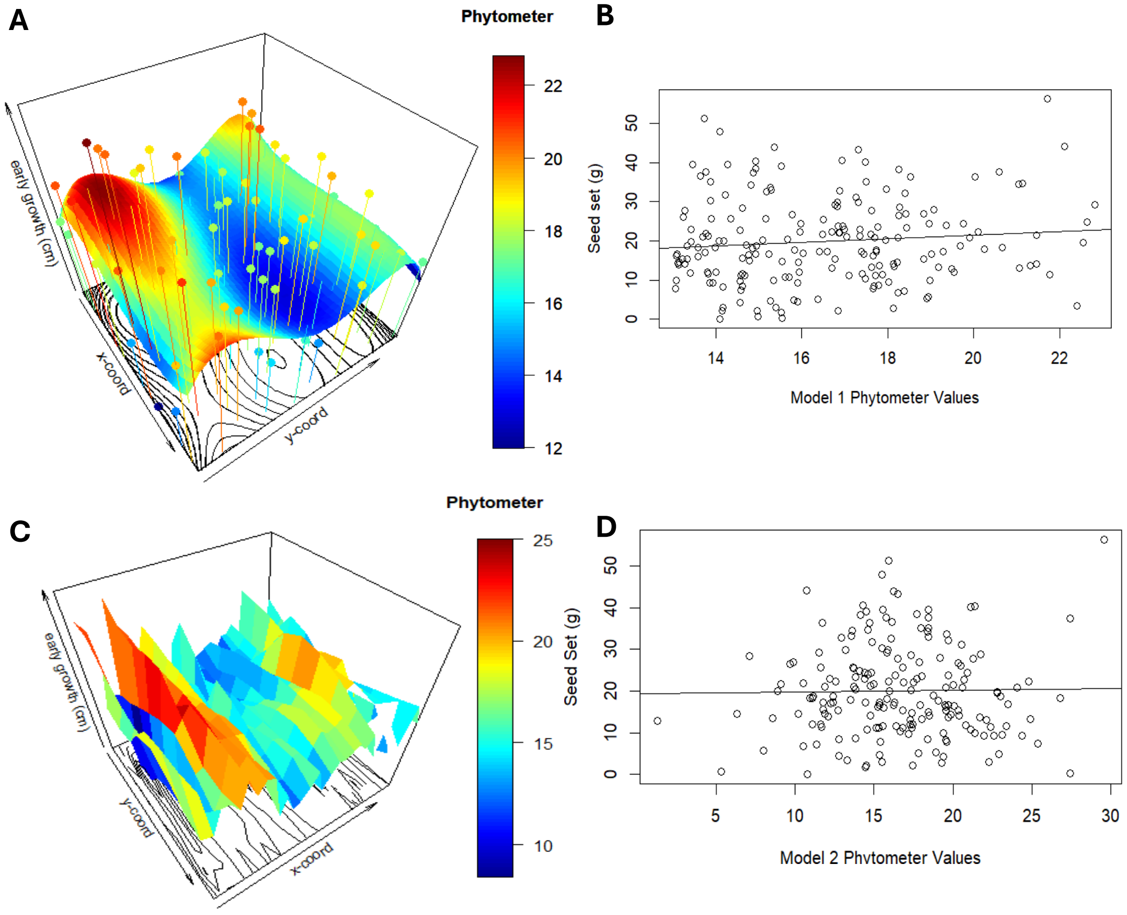


**Figure S8. Phytometer estimates do not predict seed set for a polynomial (A, B) or spline based (C,D) surface regression. A, C)** Output of polynomial (A) and spline (C) surface function for environmental quality across the four sampled plots. The 0,0 point of the xy plane is NE corner of sampled plots. Distance is in feet south (x-coord) and west (y-coord) from this point. Surface color reflects estimated surface values correlating with environmental quality, with points as *I. lacunosa* early growth values that fit the surface. As spline estimates equate to each point, they are excluded for simplicity. **B, D)** Regression estimates for both Model 1 (B) and Model 2 (D) on seed set. Neither is significant ($p=0.250$ for Model 1, $p=0.873$ for Model 2)

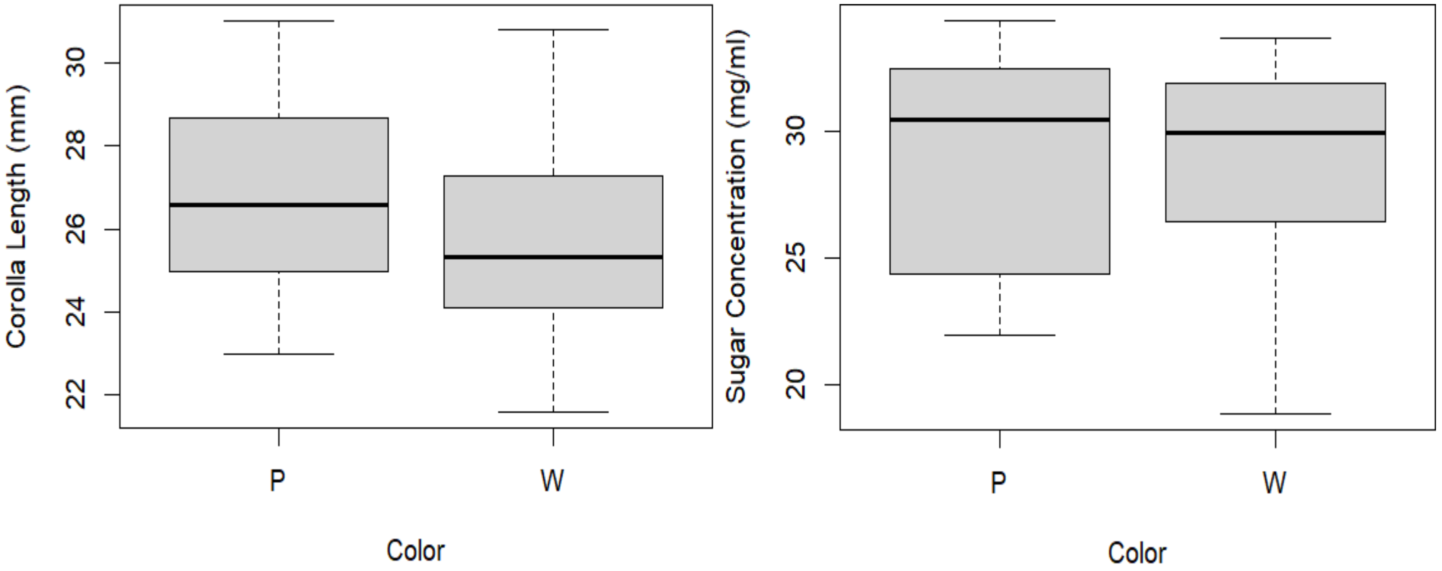


**Figure S9. Color Not Associated with Flower Size nor Sugar Concentration**. The effect of color on corolla length (left) and sugar concentration (right) in sampled RILs. Neither flower size (p-value=0.12) nor sugar concentration (p-value=0.95) significantly associates with color.

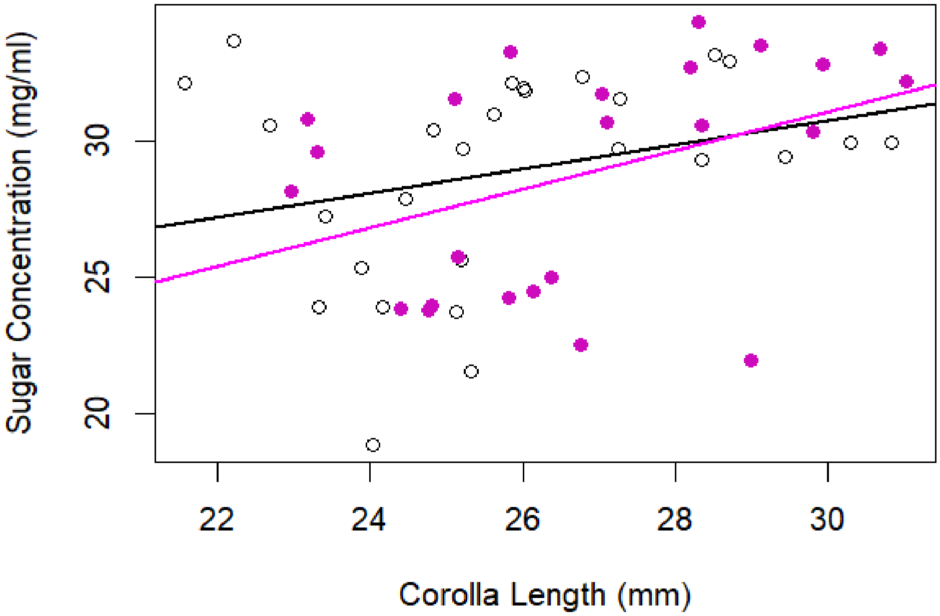


**Figure S10. Flower Size Weakly Correlates with Sugar Concentration**. The relationship between flower size and sugar concentration for pink-limbed (pink points) and white-limbed (white points) lines. While there is a significant relationship (p-value=0.018), the association is fairly weak (Adj $R^2$=0.0916) and does not differ by color.

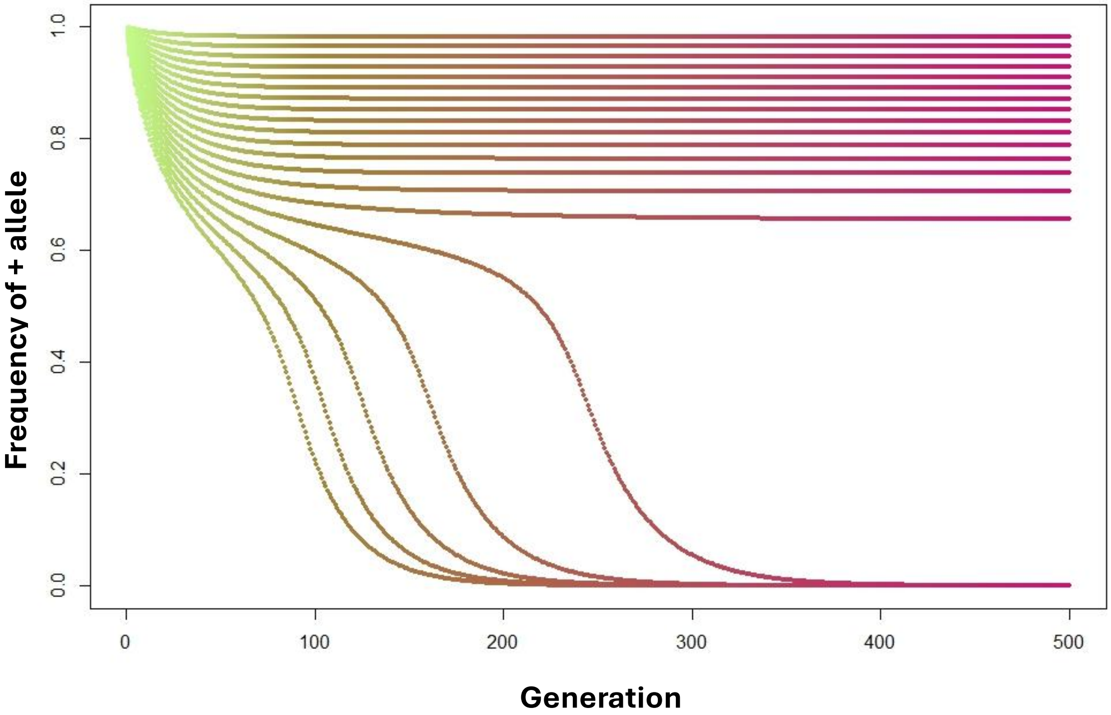


**Figure S11. Size alleles resist introgression when recurrent migration <0.032**. The change in the frequency of "+" allele at each quantitative trait locus over 500 generations given continual migration of the "-" allele haplotype. Note that all trait loci behave similarly. Migration increases by 0.02 for each trajectory. Note that the lines with migration rate greater than 0.032 (0.034, 0.036, 0.038, 0.04) result in fixation of the invading alleles, while all other lines reach a migration-selection equilibrium.

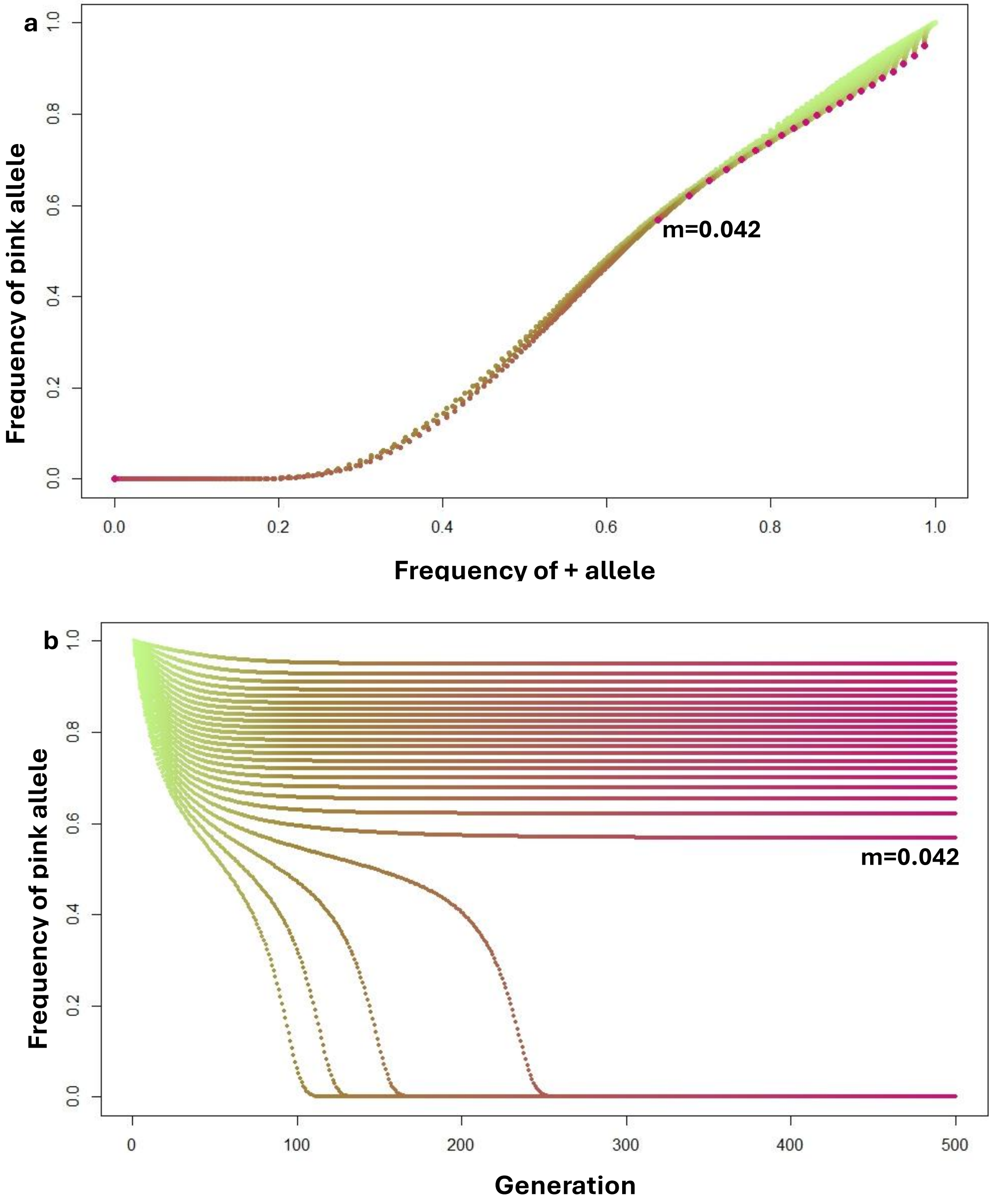
a
Frequency of pink allele
Frequency of + allele
m=0.042
0.0
0.2
0.4
0.6
0.8
1.0
b
Frequency of pink allele
Generation
m=0.042
0
100
200
300
400
500

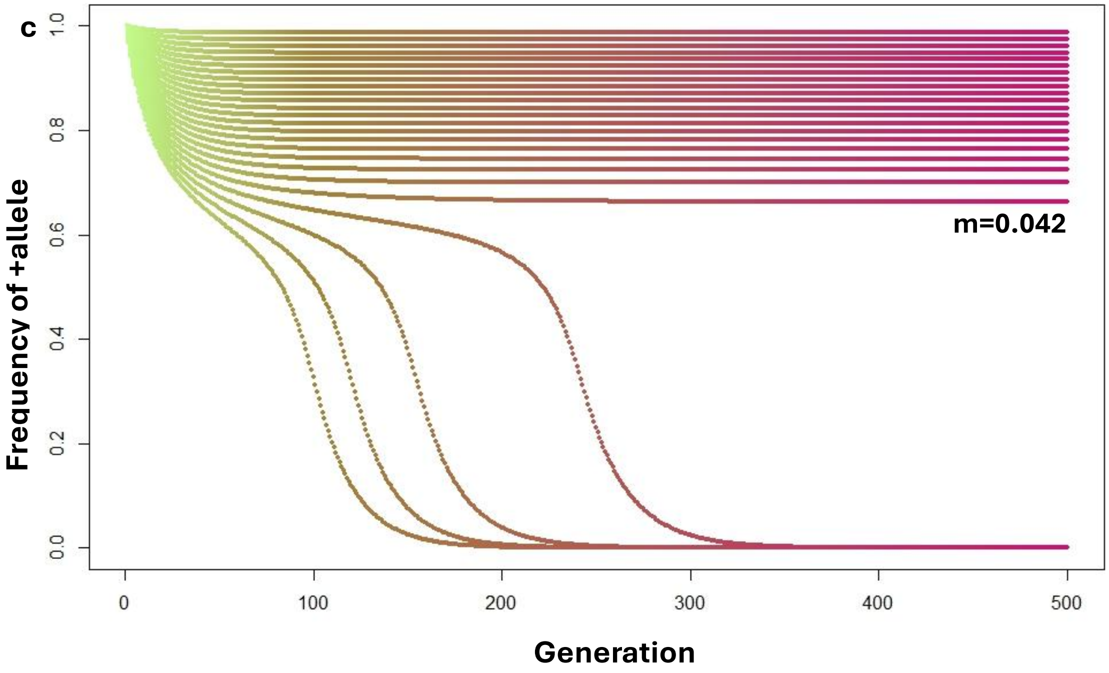


**Figure S12. Modelling joint evolution of flower color and nectary sugar concentration via total seed mass.** Parameters to generate this model is generated from data from Figure 3B and are as follows: c=$W_W$=15.6; $\beta_p$ = 1.177; $\delta z$ = 1.552667;$z_c$= 26.457. **a)** Trajectory of "pink" allele and + (*I.* cordata) sugar alleles given different rates of migration. Trajectories start with fixation of *I. cordatotriloba* alleles and end at red points. Migration rate (*m*) corresponding to the top trajectory is 0.002 and increases by 0.002 for each subsequent lower trajectory. All migration rates greater than 0.042 result in fixation of the introgressing (*I. lacunosa*) haplotype, while lower values allow color and size alleles to resist introgression via correlational selection. **b-c)** The change in the frequency of the "pink" allele (**b**) and "+" allele (**c**) over 500 generations given continual gene flow from *I. lacunosa* into *I. cordatotriloba*. Migration rates as in part A.

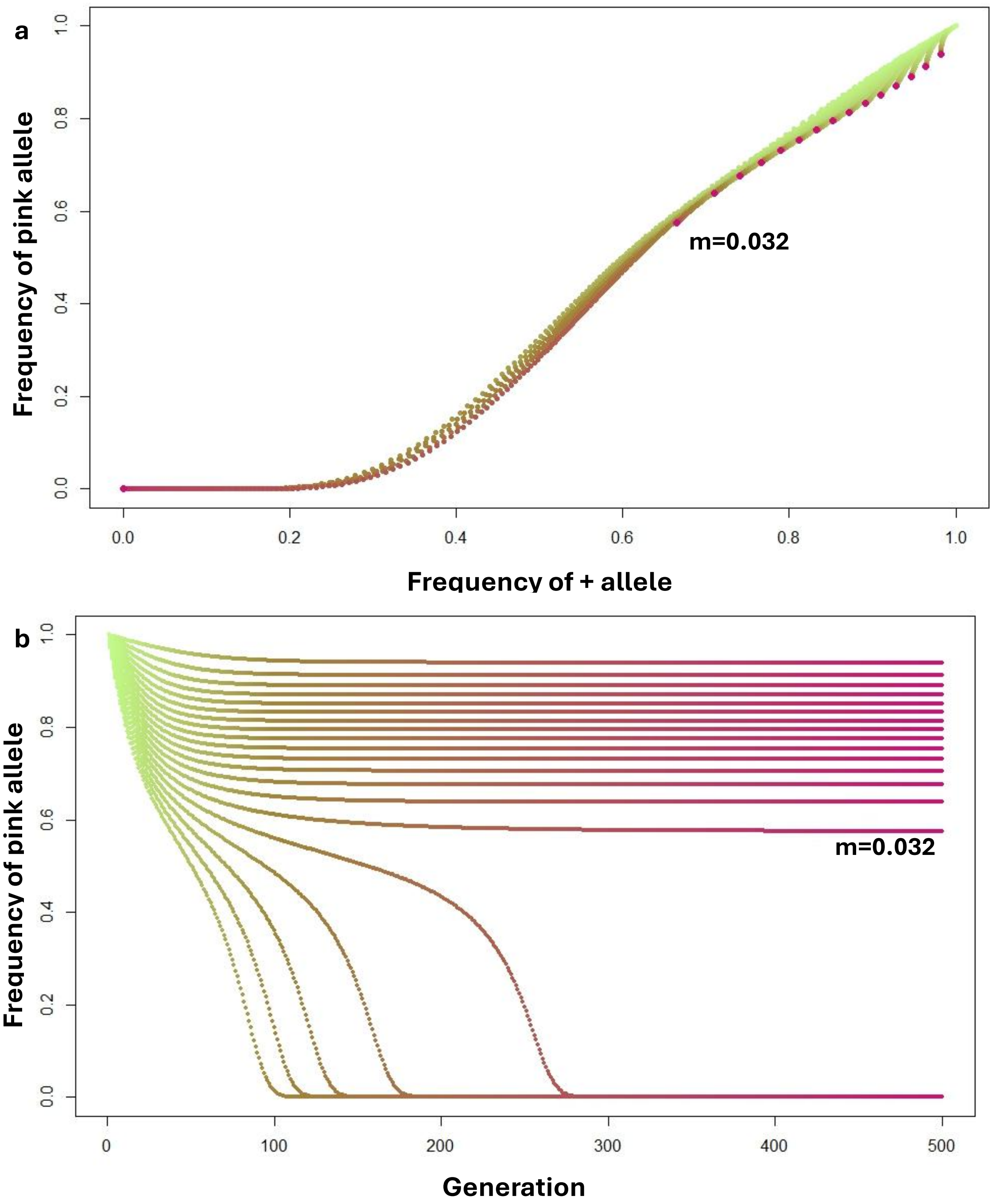
a
Frequency of pink allele
0.0
0.2
0.4
0.6
0.8
1.0
m=0.032
0.0
0.2
0.4
0.6
0.8
1.0
Frequency of + allele
b
Frequency of pink allele
0.0
0.2
0.4
0.6
0.8
1.0
m=0.032
0
100
200
300
400
500
Generation

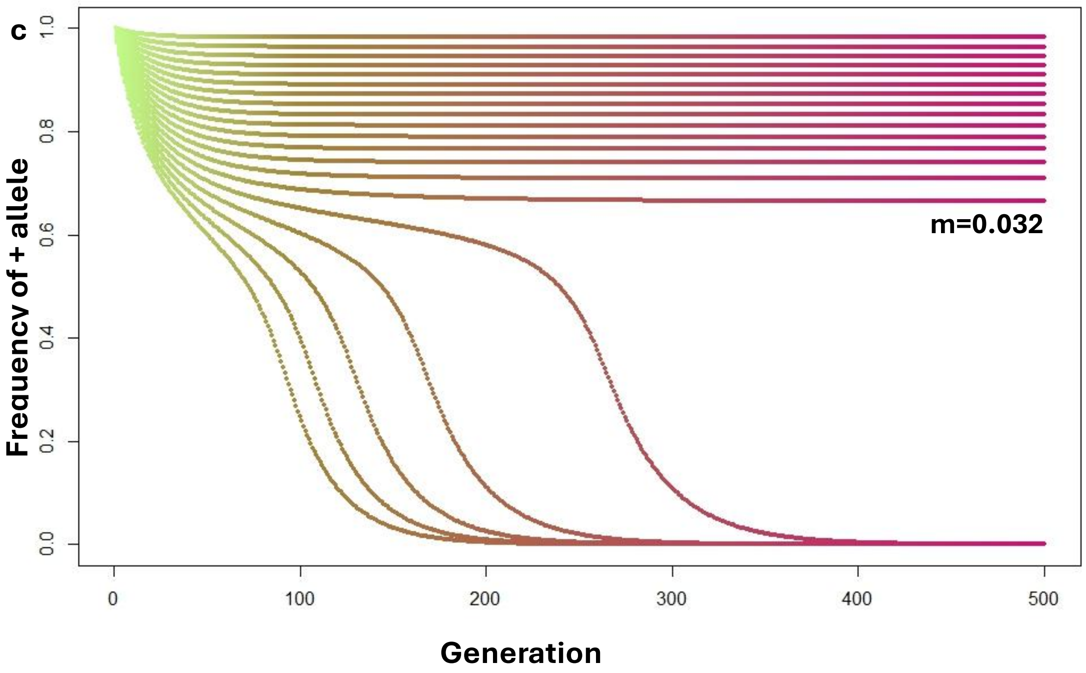


**Figure S13. Modelling joint evolution of flower color and flower size via estimated seed number.** Parameters to generate this model is generated from data from Figure 3C are as follows: $c=W_W= 957.4$; $\beta_p = 88.79$; $\delta z = 0.94755$; $z_c= 26.3$; **a)** Trajectory of "pink" allele and + (*I.* cordata) size alleles given different rates of migration. Note that because all quantitative trait loci are equivalent, the trajectory is the same at all loci. Trajectories start with fixation of *I. cordatotriloba* alleles and end at red points. Migration rate (*m*) corresponding to the top trajectory is 0.002 and increases by 0.002 for each subsequent lower trajectory. All migration rates greater than 0.03 result in fixation of the introgressing (*I. lacunosa*) haplotype, while lower values allow color and size alleles to resist introgression via correlational selection. **b-c)** The change in the frequency of the "pink" allele (**b**) and "+" allele (**c**) over 500 generations given continual gene flow from *I. lacunosa* into *I. cordatotriloba*. Migration rates as in part A.

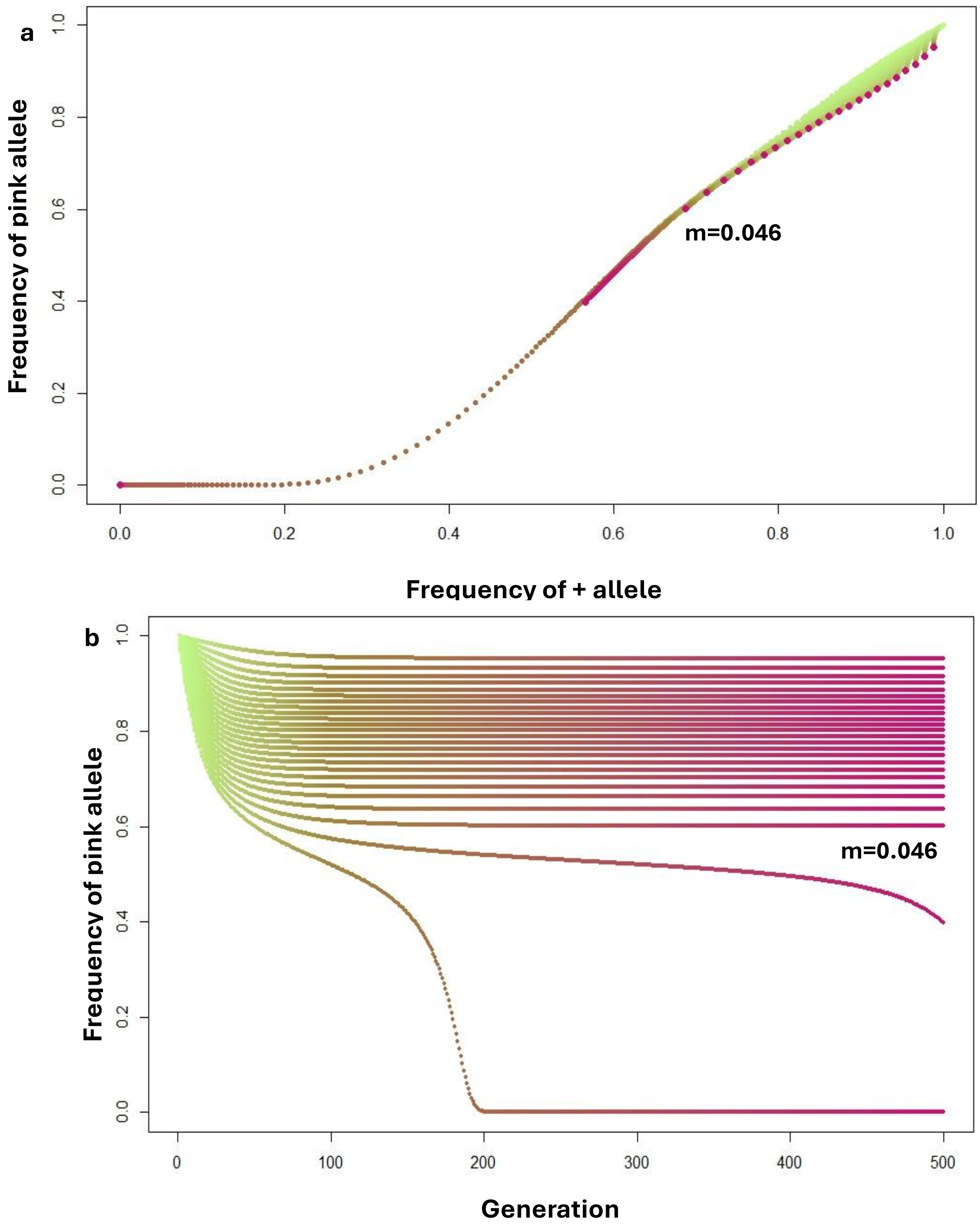
a
Frequency of pink allele
m=0.046
Frequency of + allele
b
Frequency of pink allele
m=0.046
Generation

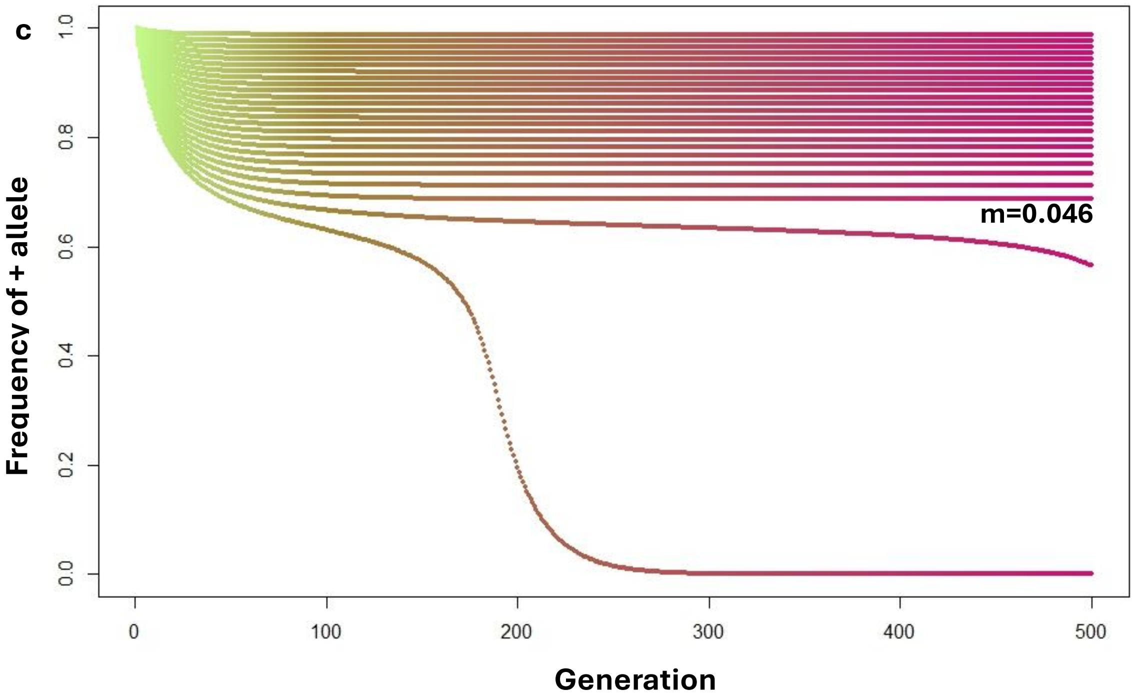


**Figure S14. Modelling joint evolution of flower color and nectary sugar concentration via estimated seed number.** Parameters to generate this model is generated from data from Figure 3D and are as follows: $W_W$=771.105; $\beta_p$ =64.29; $\delta z$= 1.55267; $z_c$=26.457. **a)** Trajectory of "pink" allele and + (*I.* cordata) sugar alleles given different rates of migration. Note that because all quantitative trait loci are equivalent, the trajectory is the same at all loci. Trajectories start with fixation of *I. cordatotriloba* alleles and end at red points. Migration rate (*m*) corresponding to the top trajectory is 0.002 and increases by 0.002 for each subsequent lower trajectory. All migration rates greater than 0.046 result in fixation of the introgressing (*I. lacunosa*) haplotype, while lower values allow color and size alleles to resist introgression via correlational selection. **b-c)** The change in the frequency of the "pink" allele (**b**) and "+" allele (**c**) over 500 generations given continual gene flow from *I. lacunosa* into *I. cordatotriloba*. Migration rates as in part A.

**Table S1. Effect of color on proportion of seed mass comprised of inviable seeds.**
Degree of freedom is 1

| | *SS* | *F* | *p* |
|---|---|---|---|
| *Color* | 0.00007 | 0.0028 | 0.9582 |

**Table S2. RADseq sites where pink-limbed allele frequencies were >0.9 for one allele in pin-limbed lines (termed pink major/white minor allele here) and >0.9 for an alternate allele in white-limbed lines (termed pink minor/white major allele here).** Haplotypes sequenced refer to both copies for a given line; number of lines is equivalent to the number of haplotypes divided by 2. Highlighted row is the chosen marker allele, as it is the only one with 24 pink lines sequenced.

| *Site* | *White RIL Haplotypes Sequenced* | *White Major Allele + Frequency* | *White Minor Allele + Frequency* | *Pink RIL Haplotypes Sequenced* | *Pink Minor Allele + Frequency* | *Pink Major Allele + Frequency* |
|---|---|---|---|---|---|---|
| *CHR6: 4796409* | 56 | A: 0.928571 | T: 0.071429 | 46 | A: 0.086956 | T: 0.913043 |
| *CHR6: 4921475* | 38 | C: 0.947368 | A: 0.052632 | 22 | C: 0.090909 | A: 0.909091 |
| *CHR6: 6720864* | 48 | T: 0.979167 | C: 0.020833 | 32 | T: 0.0625 | C: 0.9375 |
| *CHR6: 13663753* | 52 | C: 0.961538 | T: 0.038462 | 40 | C:0.05 | T:0.95 |
| *CHR6: 13740622* | 56 | T: 0.964286 | C: 0.035714 | 46 | T: 0.043478 | C: 0.956522 |
| *CHR6: 13740636* | 56 | C: 0.964286 | G: 0.035714 | 46 | C: 0.043478 | G:0.956522 |
| *CHR6: 13740637* | 56 | A: 0.964286 | T: 0.035714 | 46 | A: 0.043478 | T: 0.956522 |
| *CHR6: 15210053* | 56 | G: 0.964286 | A: 0.035714 | 48 | G:0.083333 | A: 0.916667 |
| *CHR6: 16892557* | 34 | T: 0.941176 | G: 0.058824 | 20 | T:0.1 | G:0.9 |
| *CHR6: 16892576* | 34 | G: 0.941176 | A: 0.058824 | 20 | G:0.1 | A:0.9 |

| | | | | | | |
|---|---|---|---|---|---|---|
| *CHR8: 11600238* | 2 | A:0 | G:1 | 2 | G:0 | A:1 |

**Table S3. Chi-squared tests comparing selected seeds to seeds extracted for DNA at the level of flower color or line.** All Chi-squared tests between pink and white assume a degree of freedom of 1.

| | *Color* | | *Line* | |
|---|---|---|---|---|
| | $\chi^2$ | p-value | $\chi^2$ | p-value |
| *All plots* | 0.14849 | 0.7 | 12.257, df = 47 | 1 |
| *Plot A* | 0.37875 | 0.5383 | 9.6025, df = 41 | 1 |
| *Plot B* | 0.0025866 | 0.9594 | 12.721, df = 42 | 1 |
| *Plot E* | 0 | 1 | 22.821, df = 46 | 0.9983 |

**Table S4. Numbers of plants surviving to first flower**

| | *Alive* | *Dead* | *Proportion Alive* |
|---|---|---|---|
| *Pink RILs* | 379 | 21 | 0.9475 |
| *White RILs* | 379 | 19 | 0.9523 |
| *I. lacunosa* | 398 | 3 | 0.9925 |

**Table S5. Phytometer values do not predict seed set, while early growth does.** ANOVA summary table on the effect of each predictor (early growth, phytometer values from each model) on outcome variables (seed set and early growth). Degree of freedom is 1 for all measurements.

| *Predictor* | *Outcome* | $\chi^2$ | *p-value* |
|---|---|---|---|
| *Early growth* | seed set | **17.512** | **$2.855*10^{-5}$** |
| *Model 1 values* | seed set | 1.3262 | 0.2495 |
| | early growth | **24.323** | **$8.144*10^{-7}$** |
| *Model 2-values* | seed set | 0.0257 | 0.8727 |
| | early growth | 0.1722 | 0.6782 |

**Table S6. Color does not differ between samples, even when modelling for early growth.** ANOVA summary table testing if color causes adjusted seed mass for both models to differ. Degrees of freedom is 1 unless otherwise noted.

| *Adjusted Seed Set by* | $\chi^2$ | $p$ |
|---|---|---|
| *Model 1* | 0.6145 | 0.4331 |
| *Model 2* | 2.4766 | 0.1156 |
