## Supplementary Materials and Methods for "Flower color locus resists introgression due to correlational selection with other floral traits in *Ipomoea cordatotriloba*"

***Phytometer Analysis:***

We first modeled an early growth surface based on fourth-order predictors of the phytometer row (x-coordinates) and column (y- coordinates) locations for the four plots where seeds were counted (Model 1). Using the step function in base R, we added or removed linear, quadratic, cubic, and quartic predictors to find the surface that best fit early size of *I. lacunosa*. The best fitting predictive model (lowest AIC) was as follows:

$\text{Size} = 13.082 + 0.543x + 1.757y - 0.0148x^2 - 0.189y^2 + 0.000102x^3 + 0.00662y^3 - 0.0000722y^4 - 0.0180xy + 0.000262x^2y - 3.166*10^{-6}x^3y$.

Because a polynomial surface such as that fitted using Model 1 may not accurately model surfaces with multiple peaks or with sharp changes in slopes, we additionally modeled the phytometer surface using a spline-based approach (Model 2) via the MBA package in R (Finley et al. 2024). Planar splines were created by drawing planes between the *I. lacunosa* early growth measurements within the four plots surveyed. For both Models 1 and 2, we used the phytometer surface to calculate an adjusted seed mass for individual *I. cordatotriloba* plants by dividing observed seed mass by the estimated surface values of the predictive model to account for differences in seed set ascribable to local micro-environmental variation. All models were visualized using the plot3D library in RStudio (Soetaert and Soetaert 2024).

***Likelihood Model for Siring Success:***

Here we describe the likelihood model we used to estimate siring success of plants with pink- and white-limbed flowers. Maternal and offspring genotypes are scored for genotype at a SNP with alleles “A” and “G”. 22 RILs with pink limbs are “AA”, one is “GG”, and none are “AG”. By contrast, 26 RILs with white limbs are “GG”, one is “AG” and none are “AA”.

The basic data is the number of sampled offspring for each combination of maternal and offspring genotypes at the SNP. Let the number of offspring of genotype j produced by pink-limbed maternal plants of genotype i be $pN_{i \to j}$ (i, j ∈ {AA, AG, GG}. The corresponding number for white maternal plants is $WN_{i \to j}$ . Then the Likelihood of the data is

$$\boldsymbol{L} = C \prod_{i,j} {}^{P}P_{i \to j}^{\;{}^{P}N_{i \to j}} \cdot \prod_{i,j} {}^{W}P_{i \to j}^{\;{}^{W}N_{i \to j}} \qquad \text{Eq. (4)}$$

where ${}^{P}P_{i \to j}$ is the probability that a pink-limbed maternal plant of genotype i produces an offspring of genotype j, and ${}^{W}P_{i \to j}$ is the corresponding probability for white -limbed maternal plant. C is a combinatorial constant eliminated in the analysis. The expressions for ${}^{P}P_{i \to j}$ and ${}^{W}P_{i \to j}$ are given in Table SA1, with the variables defined in Table SA2.

To illustrate how these probabilities are obtained, I derive here the probability that a pink-limbed maternal parent of genotype AA produces an offspring of genotype AA, ${}^{P}P_{AA \to AA}$ ; all other probabilities can be derived following the same process. For an AA maternal parent, an AA offspring can either occur through selfing, with probability sp, or through outcrossing with an A pollen grain, with probability 1 – sp. If it selfs, the probability that the offspring is AA is 1, whereas if it outcrosses, that probability is P(A), the probability outcross pollen carries allele A:

$${}^{P}P_{AA \to AA} = s_P + (1- s_P)*\mathrm{P(A)}$$

A-carrying pollen can be produced either by pink-limbed or white-limbed plants. The probability pollen is from a pink-limbed plant and carries A is Pa, the probability it is from a pink-limbed plant times the probability that such pollen carries A. Analogously, the probability that pollen is from a white-limbed plant and carries A is $(1-g)(1-p`)$, the probability that pollen from a white plant carries A times the probability that a pink -limbed plant is sired by a white-limbed plant. Thus, we have the following equation:

$${}^{P}P_{AA \to AA} = s_P + (1- s_P)\,(ap` + (1-g)(1-p`)).$$

Substituting the values in Table SA1 into Eq. (4) yields

$$\boldsymbol{L} = C * \left[s_P \ + (1- \ s_P)\big(ap` + (1-g)(1-p`)\big)\right]^{{}^{P}N_{AA \to AA}}$$
$$* \ ((1- \ s_P)((-a)p` \ g(1-p`))^{{}^{P}N_{AA \to AG}}$$
$$* \ \big((1- \ s_P)(ap` + (1-g)(1-p`))\big)^{{}^{P}N_{GG \to AG}}$$
$$* \ \big(s_P + (1- \ s_P)((1-a)p` + g(1-p`)\big)^{{}^{P}N_{GG \to GG}}$$
$$* \ (s_w \ + (1- \ s_w)((1-g)w` + (1-a)w`))^{{}^{W}N_{AA \to AA}}$$
$$* \ \big((1- \ s_w)(gw` + (1-a)(1-w`))\big)^{{}^{W}N_{AA \to AG}}$$
$$* ((s_w)/4 \ + (1- \ s_w)((1-g)w` + a(1-w`))/2)^{{}^{W}N_{AG \to AA}}$$
$$* \ (1/2)^{{}^{W}N_{AG \to AG}}$$

$$* ((s_w)/4 + (1 - s_w)(gw` + (1 - a)(1 - w`))/2)^{wN_{AG \to GG}}$$
$$* ((1 - s_w)((1 - g)w` + (1 - a)w`))^{wN_{GG \to AG}}$$
$$* (s_w + (1 - s_w)(gw` + (1 - a)(1 - w`)))^{wN_{GG \to GG}}$$

To estimate these parameters, we performed a log-likelihood optimization for 300 randomized starting conditions to prevent false parameter estimates on local but not global maxima. We then recorded the values at these six parameters and a log-likelihood score. Once we calculated values for the unconstrained model, we then tested if constrained models are significantly worse predictors than the unconstrained model via log-likelihood ratio tests. We constructed three models that included different sets of constraints. The first (Model 1) fixed *a* and *g* to their known values – the frequency of the A allele in sampled pink-limbed lines (*a*=0.9565) and the frequency of the G allele in sampled white-limbed lines (*g*=0.9821). The second model (Model 2) built upon Model 1's constraint (a=0.9565; *g*=0.9821) and added the constraint that selfing rates were equal regardless of flower color ($s_P = s_W$). The third model (Model 3) built upon the prior two models by assuming that the rate of per-capita white-limbed pollen siring white-limbed plants is the same as the rate of pink-limbed pollen siring pink-limbed plants (W=P).

Model testing was conducted sequentially. We first compared model 1 with model 0. If model 1 was significantly worse than model 0, we would test model 2 against model 0; if it was not significantly worse, we would test model 2 against model 1. Finally, if model 2 was significantly worse than the model it is compared to (model X), we would test model 3 against model X. Otherwise, we would test model 3 against model 2. To test whether W = P, and thus whether pink- and white-limbed plants have equal *per-capita* siring success, we asked whether Model 3 produces a significantly lower likelihood estimate than the model with the most constraint not significantly different from the unconstrained model.

Table A.1. Probabilities of offspring genotype for a given maternal genotype. Probabilities are listed separately for pink- and white-limbed plants.

A. Probabilities (${}^{P}P_{i \to j}$) for pink-limbed maternal parents:

| | Maternal Genotype | |
|---|---|---|
| Offspring Genotype | $AA$ | $GG$ |
| $AA$ | $s_P + (1 - s_P)(ap\grave{} + (1 - B)(1 - p\grave{}))$ | 0 |
| $AG$ | $(1 - s_P)((1 - a)p\grave{} + B(1 - p\grave{}))$ | $(1 - s_P)(ap\grave{} + (1 - B)(1 - p\grave{}))$ |
| $GG$ | 0 | $s_P + (1 - s_P)((1 - a)p\grave{} + B(1 - p\grave{}))$ |

B. Probabilities ($^{W}P_{i\to j}$) for white-limbed maternal parents

| | Maternal Genotype | | |
|---|---|---|---|
| Offspring Genotype | $AA$ | $AG$ | $GG$ |
| $AA$ | $s_w + (1- s_w)((1-g)w\grave{} + a(1-w\grave{}))$ | $(s_w)/4 + (1- s_w)((1-g)w\grave{} + a(1-w\grave{}))/2$ | 0 |
| $AG$ | $(1- s_w)(gw\grave{} + (1-a)(1-w\grave{}))$ | 1/2 | $(1- s_w)((1-g)w\grave{} + a(1-w\grave{}))$ |
| $GG$ | 0 | $(s_w)/4 + (1- s_w)(gw\grave{} + (1-a)(1-w\grave{}))/2$ | $s_w + (1- s_w)(gw\grave{} + (1-a)(1-w\grave{}))$ |

Table A2. Definition of parameters and variables.

| Term | Definition |
|---|---|
| $s_p$ | Selfing rate of pink-limbed plants |
| $s_w$ | Selfing rate of white-limbed plants |
| $a$ | probability that pollen from a pink-limbed plant is $A$ |
| $g$ | probability that pollen from a white-limbed plant is $G$ |
| $p$` | probability a pink-limbed plant is pollinated by pollen from a pink-limbed plant |
| $w$` | probability a white-limbed plant is pollinated by pollen from a white-limbed plant |

***Calculating Frequency Dependence of Per-capita Siring Success***

Equations (2a, b) give the per-capita siring success of pink- and white-limbed plants. This success is inherently frequency dependent, as can be seen from the following considerations: Letting $N_p$ and $N_W$ be the numbers of pink- and white-limbed plants in the population, respectively, then the proportion that are pink is:

$$f = \frac{N_p}{N_p+N_w} = \frac{N_p}{N}, \text{ where } N = N_p + N_W .$$

Equations 2a and 2b can thus be re-written as

$$F_P^* = \frac{f\,N\,(s+(1-s)p)+(1-f)N(1-s)(1-w)}{f\,N} \qquad \text{Eq. (5a)}$$

and

$$F_W^* = \frac{(1-f)N\,(s+(1-s)w)+fN(1-s)(1-p)}{f\,N} \qquad \text{Eq. (5b)}$$

assuming $s_P = s_W = s$ as before. Because the $N$'s cancel, these reduce to

$$F_P^* = s + (1-s)P + \frac{1-f}{f}\,(1-s)(1-W) \qquad \text{Eq. (6a)}$$

and

$$F_W^* = \frac{1-f}{f}\,(s + (1-s)W) + (1-s)(1-P) \qquad \text{Eq. (6b)}$$

Because these are functions of $f$, fitnesses $F_P^*$ and $F_W^*$ are inherently frequency dependent.

In our experiment, $f = ½$, so $\frac{1-f}{f} = 1$ and Eqs. (6a, 6b) reduce to

$$F_P^* = (s + (1-s)P) + \ (1-s)(1-W) \qquad \text{Eq. (7a)}$$

$$F_W^* = \ (s + (1-s)W) + (1-s)(1-P) \qquad \text{Eq. (7b)}$$

From these equations, it can be seen that $F_P^* = F_W^*$ implies and is implied by $P = W$. Thus, testing whether $P = W$ provides a test of whether pink- and white-limbed plants have equal siring success.

***Modelling Joint Selection and Recurrent Introgression - Mathematica Model:***

This model built in SAS specifies color phenotype to be determined by genotype at a single "color" locus, while the value of the quantitative trait is determined additively by the effects of two alleles at each of $k$ unlinked loci. One allele, the "+" allele, increases the trait by a fixed increment, $\delta z$, while the "–" allele decreases the trait by the same fixed increment (Figure 1). An individual gamete type $i$ is represented by a vector, $\boldsymbol{g}_i$, with $k + 1$ elements, all either 1 or 0. If the first element is 1, the gamete carries the "white" allele, while if if it is 0, it carries the "pink" allele. In the subsequent elements, 0 indicates a "+" allele while 1 indicates a "–" allele. The ordering of gametes is given by $\boldsymbol{g}_i$ = Binary[$i$], which is $i$ expressed as a base 2 number with enough preceding 0's to yield $k$+1 elements. For example, with $k = 5$, Binary[5] = { 0, 0, 0, 1, 0, 1}.

With $n$ loci, there are $n_g = 2^n$ possible gametes. The gamete frequencies at time $t$ are represented by the vector $\boldsymbol{x^t} = \left\{x_1, x_2, \cdots, x_{n_g}\right\}$. The program implementing this model has the following steps.

<u>1. Set initial gamete frequencies</u>:

$$x^0 = \{1, 0, 0, \cdots, 0\}$$

This represents fixation of the gamete with the pink allele at the color locus and "+" alleles at all quantitative trait loci.

<u>2. Implement immigration.</u>

$$x^{0'} = (1-m)x^0 + m\,\{1, 1, 1, \cdots,\} \ ,$$

Where $m$ is the proportion of immigrant gametes after immigration.

<u>3. Calculate genotype frequencies</u>

Let $\mathbf{G_{ij}}$ designate the genotype produced by gametes $\boldsymbol{g_i}$ from parent 1 and $\boldsymbol{g_j}$ from parent 2. Genotype frequencies are represented by an $n_g$ x $n_g$ matrix $\boldsymbol{X}$, where the element $X_{ij}$ is the frequency of genotype $\mathbf{G_{ij}}$ . This matrix is calculated as

$$\boldsymbol{X} = x^{0'} \otimes \; x^{0'}$$

where $\otimes$ indicates outer product. It assumes mating is random.

4. Impose selection

Fitnesses are represented by an $n_g$ x $n_g$ matrix, $\boldsymbol{W}$, in which element $W_{ij}$ is the fitness of genotype $\mathbf{G_{ij}}$ . $W_{ij}$ is determined by the number of +" and "–" alleles that genotyhpe carries. Specifically

$$W_{ij} = \begin{cases} \alpha + \beta\, \delta z\, (n_+ - n_-) \text{ if the genotype carries at least 1 pink allele} \\ K \text{ if the genotype carries no pink alleles} \end{cases}$$

where $n_+$ and $n_-$ are the numbers of "+" and "–" alleles, respectively carried by the genotype and $\alpha$ is the fitness of a genotype with an equal number "+" and "– " alleles, and $\delta z$ is the change in mean phenotype resulting for a "–" allele to a "+" allele.

After selection, the new genotype frequencies are

$$\boldsymbol{X'} = \frac{\boldsymbol{X}\,\boldsymbol{W}}{\sum_i \sum_j X_{ij} W_{ij}}$$

5. Calculate new gamete frequencies

New gamete frequencies, $\boldsymbol{x^1}$, are generated using a reproduction tensor, $\boldsymbol{T_R}$, which is of rank 3, with each dimension having $n_g$ elements. The element $(T_R)_{ij}$ is a vector of gamete frequencies, analogous to $\boldsymbol{x}$, produced by genotype $\mathbf{G_{ij}}$ .

The new gamete frequencies are generated by

$$\boldsymbol{x^1} = \sum_i \sum_j X_{ij} (T_R)_{ij}$$

After this step is completed, return to step 2 for the next generation.

Allele frequencies were calculated at the beginning of each generation. We calculated the frequency of the pink allele as $\sum_i x_i$ where $i$ is the set of gametes with the pink allele (first element of gamete is 0). We calculated the frequency of the "+" at quantitative trait locus $j$ as $\sum_k x_k$ where $k$ is the set of gametes with a "+" allele at locus $j$.

The mean value of the quantitative trait, $\bar{z}$ , was also calculated each generation. To do so, we initially established a phenotype matrix, $\boldsymbol{Z}$. Each element $Z_{ij}$ of this matrix is the trait value for genotype $\mathbf{G_{ij}}$ and was calculated as

$$Z_{ij} = M\ \delta z\ (n_+ - n_-)$$

Mean phenotype was then calculated as

$$\bar{z} = \sum_i \sum_j X_{ij} Z_{ij}$$

The Mathematica program used to implement this model is presented below and will be deposited in an appropriate repository upon acceptance of the manuscript.

***Modelling Joint Selection and Recurrent Introgression - R Model:***

The model built in R follows the same initial set up as the one built in SAS; it specifies color phenotype to be determined by genotype at a single "color" locus, while the value of the quantitative trait is determined additively by the effects of two alleles at each of $k$ unlinked loci. One allele, the "+" allele, increases the trait by a fixed increment, δz, while the "–" allele decreases the trait by the same fixed increment (Figure 1). An individual gamete type $i$ is represented by a vector, $\boldsymbol{g}_i$, with $k + 1$ elements, with values of the first element represented by either 1 for carrying the "pink" allele or 0 for the "white" allele, and later elements represented by 1 for a "+" allele and 0 for a "–" allele. The ordering of gametes is given by $\boldsymbol{g}_i$ = Binary[$i$], which is $i$ expressed as a base 2 number with enough preceding 0's to yield $k$+1 elements. We likewise run this model in R in five stages.

(1) Set initial gamete frequencies:

We designate each of the $2^{k+1}$ possible gametes by $g_i$, $i = 1 ,2, . . . 2(k+1)$, letting $g_1$ be the genotype with all "–" alleles for the quantitative trait and with the "white" allele at the flower-color locus. We then further designated the gamete frequencies as $\boldsymbol{x} = \{x_1, x_2, . . . , x_{2(k+1)}\}$, where $x_i$ is the frequency of gamete $g_i$.

(2) Implement immigration:

Starting with gamete frequencies $\boldsymbol{x}^t$ at time $t$, a pulse of immigration produces a new set of gamete frequencies:

$$\boldsymbol{x}^{t'} = m\,\{1, 0, 0, \cdots, 0\} + (1 - m)\boldsymbol{x}^{\,t}$$

where ***m*** is the fraction of all gametes that are immigrants.

(3) Calculate genotype frequencies:

Next, we consider each possible genotype, γ. That genotype can be produced by one or more combinations of gametes. Let $C_l^\gamma = \{g_{i_l}, g_{j_l}\}$ be the $l^{th}$ such combination, with involves gametes $g_{i_l}$ and $g_{j_l}$. Here $l$ runs from 1 to $n_l$, where $n_l$ is the total number of combinations of gametes. Under random mating, the frequency of the genotype produced by gamete combination $C_l^\gamma$ is

$$\pi_l^\gamma = x_{i_l}^{t\,'} x_{j_l}^{t\,'}$$

where $x_{i_l}^{t\,'}$ is the frequency of gamete $i_l$ and $x_{j_l}^{t\,'}$ is the frequency of gamete $j_l$. The total frequency of genotype γ is then

$$X^\gamma = \sum_{l=1}^{n_l} \pi_l^\gamma$$

(4) Impose selection:

The fitness of genotype γ, designated $W_\gamma$, is determined by the number of "+" and "–" alleles it carries. Specifically,

$$W_\gamma = \begin{cases} \alpha + \beta\,\delta z\,(n_+ - n_-) \text{ if the genotype carries at least 1 pink allele} \\ K \text{ if the genotype carries no pink alleles} \end{cases}$$

where $n_+$ and $n_-$ are the numbers of "+" and "–" alleles, respectively carried by the genotype and $\alpha$ is the fitness of a genotype with equal numbers of "+" and "– " alleles, and $\delta z$ is the change in mean phenotype resulting for a "–" allele to a "+" allele. After selection, the new frequency of genotype γ is

$$X^{\gamma\prime} = \frac{X^\gamma W_\gamma}{\sum_\theta X^\delta W_\theta} ,$$

where $\theta$ is the set of all genotypes.

(5) Calculate new gamete frequencies:

The new frequency of gamete *i* is calculated as

$$x_i^{t+1} = \sum_\gamma \varepsilon_\gamma$$

where

$$\varepsilon_\gamma = \begin{cases} 0 \text{ if genotype } \gamma \text{ cannot produce gamete } g_i \\ \dfrac{1}{2^\tau} \text{ if genotype } \gamma \text{ can produce gamete } g_i \end{cases}$$

and $\tau$ is the number of heterozygous loci in that genotype.

Allele frequencies were calculated at the end of each generation. As with the SAS model, the frequency of the color allele was calculated as the sum of all haplotypes with the pink color allele. Likewise, we independently calculated the frequency of the + alleles for the size locus in the same manner (sum of gamete frequencies with the + allele at each locus). Because the size alleles do not have linkage disequilibrium with each other, values of size alleles are identical across loci and a size allele is chosen at random out of the k locus to represent frequency of size loci. The R program used to implement this model is presented below and will be deposited in an appropriate repository upon acceptance of the manuscript.
